# Shuttling, swapping and mixing: the rapid modular evolution of antiviral repertoires in temperate phages and their satellites

**DOI:** 10.64898/2026.02.27.708500

**Authors:** Jorge Moura de Sousa, Florence Depardieu, Marie Touchon, Raphaël Laurenceau, Kristen Curry, Alice Maestri, Florian Tesson, Arthur Loubat, Rayan Chikhi, Jean Cury, Aude Bernheim, David Bikard, Eduardo P.C. Rocha

**Affiliations:** Institut Pasteur, Université de Paris Cité, CNRS UMR3525, Microbial Evolutionary Genomics, Paris, France; Institut Pasteur, Université de Paris Cité, CNRS UMR3525, Synthetic Biology, Paris, France; Institut Pasteur, Université de Paris Cité, CNRS UMR3525, Sequence Bioinformatics, Paris, France; Institut Pasteur, Université de Paris Cité, CNRS UMR3525, Molecular Diversity of Microbes, Paris, France

## Abstract

Interactions between bacteria, bacteriophages, and their satellites are shaped by a myriad of defence and counter-defence mechanisms. Here, we identified and characterized the defence hotspots of thousands of P2-like phages and P4-like satellites to elucidate the origins and evolutionary dynamics of defence systems. Both P4 and P2 encode a broad diversity of recognizable defence systems. Defences are a substantial, yet likely underestimated, share of the elements’ pangenomes, as shown by novel antiviral functions discovered in P4 loci lacking known defence genes. Defence loci are very rapidly swapped, without pseudogenization, suggesting defences are replaced before becoming non-adaptive. This intense local recombination melds components of distinct systems into novel functional chimeras. Systems swap so rapidly that many elements with identical core genes have completely different defences. Surprisingly, despite P4 and P2’s concomitant replication and packaging, they almost never exchange defence genes. In contrast, near identical defence systems can be found in distinct types of MGEs and in cryptic chromosomal locations. Our findings highlight P4 and P2 as mobile platforms driving the modular diversification of bacterial antiviral repertoires. Hence, bacterial defences change quickly by phage and satellite turnover, and by the quick swap of defences within these elements.

## Introduction

Microbial ecosystems are shaped by complex interactions between bacteria and bacteriophages (phages). These interactions range from the direct antagonism between virulent phages and bacteria^1^ to the more nuanced continuum between antagonism and commensalism of temperate phages and their hosts, where the interests of both bacteria and phage can be aligned^2,3^. One key example of the latter is the presence of antiviral defences in prophages, which protect their host bacteria from predation by other phages, thus ensuring the survival of both^4,5^. Prophages are not the only mobile genetic elements (MGEs) that provide antiviral functions to their hosts. Anti-MGE defence systems are frequently encoded by MGEs present in bacterial genomes^6^, possibly because of competition, and/or cooperation, between MGEs^7^. This is exemplified by phage satellites, non-autonomous mobile elements (hitchers) that hijack (helper) phages to mobilize between bacteria^8,9^. Some P4-like satellites encode antiviral defences against competitors of P2-like helper phages, protecting both helper and bacterial host^10^. Hence, the associations between satellites, phages and bacteria evolve in a mutualism-parasitism continuum, where partners incur in costs and benefits associated with the presence of the other elements^8^.

The diversity of known defence systems is rapidly increasing. Defence genes have one of the highest rates of gain/loss across bacterial genomes^11^, and in bacteria, their acquisition is typically associated with the mobility of the MGEs carrying them^12^. But little is known about the evolutionary dynamics of defence genes in MGEs themselves. Where do these genes come from, and how stable are they? What is the fate of these genes, and that of the MGEs carrying them, once defences are no longer useful? These questions are intricately linked not only with the ecology of MGEs (who is defending whom, when and against what), but also with their genomic organization, which for some elements like phages and satellites, can be tightly conserved^9,13,14^. The accumulation of accessory genes in phages is constrained by the size of their genomes, which must fit within their capsids^15^, and the very modular genomic organization of both satellites and phages^9,14^. Although prophages often have accessory genes useful to their bacterial hosts, their high gene density suggests they rarely have pseudogenes and is thought to explain the low frequency of transposable elements in their genomes, in contrast with plasmids^16^. Hence, the systematic detection of defence genes in prophages and satellites reflects the selective importance of these functions for their ecology and evolution. At a broader level, understanding the prevalence, localization and evolutionary dynamics of antiviral genes can highlight the rules that govern the genomic plasticity and genomic organization of MGEs, as well as their ecological associations and interactions with bacterial hosts and with other MGEs.

Several defence systems were previously detected in P2-like phages and P4-like satellites, in a single hotspot near the *cos* site of each element^10^. Here, we delimit satellite and phage genomes to identify additional hotspots and to account for MGE similarity when studying the evolutionary dynamics of antiviral genes within and across P4 satellites and P2 helpers. To extend the diversity of our dataset, we coupled the detection and precise delimitation of thousands of P4-like satellites^17^ and P2-like prophages in complete bacterial genomes with thousands of additional P4-like satellites detected in metagenomes to generate the most abundant and diverse genomic collection of satellites and helper phages to date. The extensive dataset of P4 and P2 genomes allowed to uncover the quick turnover of defence genes encoded by P4 and P2, the genetic exchanges that underly this turnover, and how they shape defence repertoires.

## RESULTS

### Defence systems’ loci are pervasive in a few hotspots of P2 and P4

We identified the P2-like prophages (P2 for short) and P4-like satellites (P4 for short) in the complete genomes of *Escherichia coli, Klebsiella pneumoniae, Salmonella enterica*. We also used the P4-like contigs detected in Logan, a repository of assembled sequence data from tens of millions of public sequencing experiments^18^, which greatly expanded the number and genomic diversity of known P4 satellites (Fig S1, see Methods). The 3085 P4 (from complete bacterial genomes and metagenomes) and 1511 P2 (from complete bacterial genomes) elements in our dataset were dereplicated and genetically delimited at the nucleotide level by identification of their *att* sites (see Methods). We then identified the defence systems in P4 and P2 using DefenseFinder^19^ and PADLOC^20^. We also searched for anti-defence systems, for which we only find two types (Anti_RM and Anti_CRISPR) distributed in 28 P2. Most of these P2 have a single anti-antiviral system, but one has five distinct systems. We detected no anti-defence systems in P4. Given the low numbers of these anti-defence systems, we decided to exclude them from the main analysis.

We found a very broad diversity of known types of defence systems, which occurred in 57% of P4 and in 49% of P2 (Fig 1A and B, Fig S2A for PADLOC classification). Some types of systems were detected in both satellites and phages (e.g., AbiU or PD-T7-2), whereas others were found in only one of them (e.g., AbiP2 in P2 and Hachiman in P4). We also identified individual genes that are usually associated with defence systems, but that do not form a complete known system (see the two rightmost columns in Fig 1A for systems that were partially detected by DefenseFinder). We will refer to these as putative defence systems and will explore them in more detail in a subsequent section. If we include both complete and putative systems, the proportion of elements that encode defence-associated genes increases to 73% (P4) and 81% (P2) (Fig 1B). Hence, a large majority of these two types of MGEs encode genes associated with antiviral activity.

**Figure 1.**
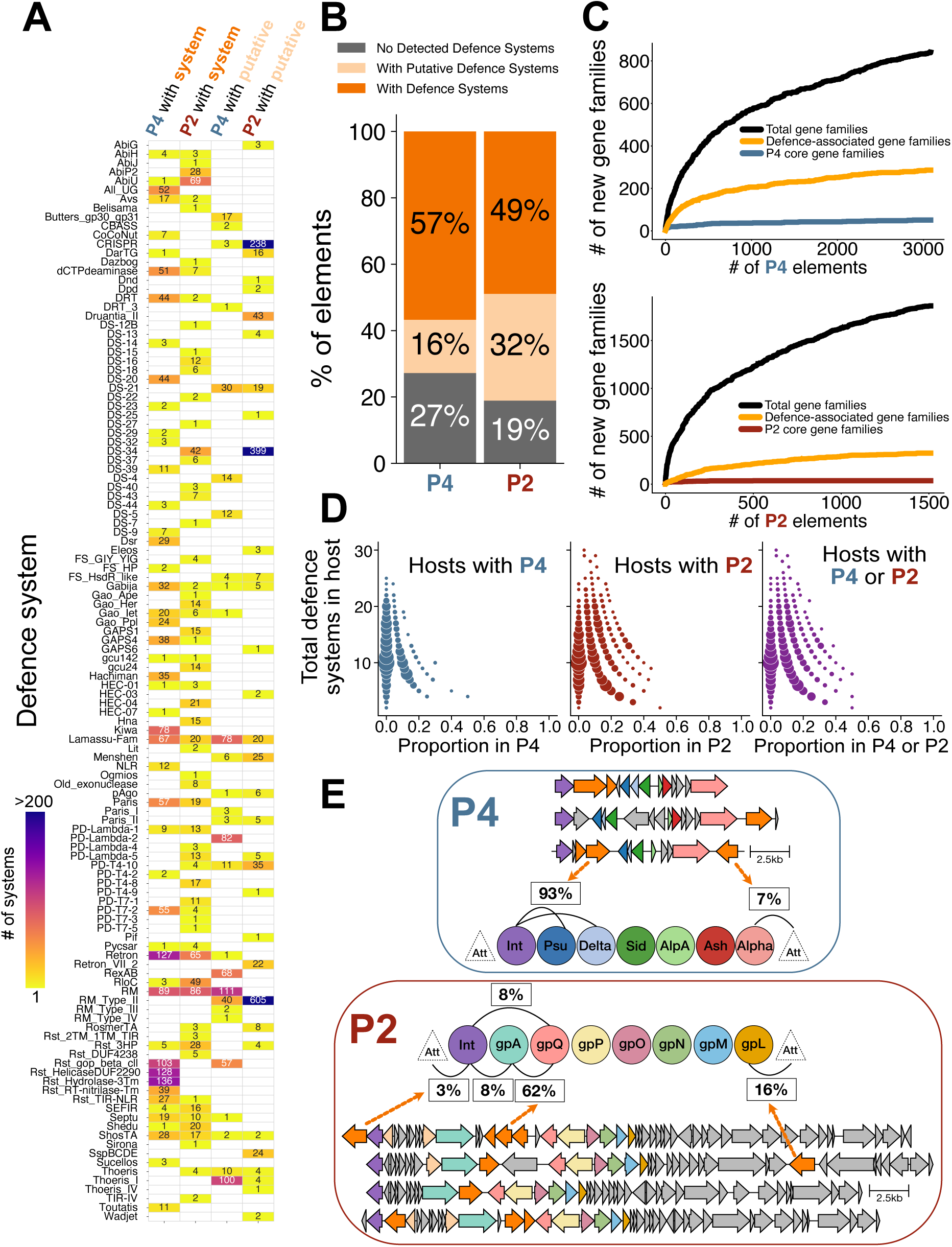
Abundance, distribution and localization of defence systems in P4-like satellites and in P2-like prophages. **A.** Number of the different types of complete (two leftmost columns) and putative (two rightmost columns) defence systems detected by DefenseFinder in P4 (1^st^ and 3^rd^ column) and P2 (2^nd^ and 4^th^ column). Darker colours in the heatmap correspond to higher numbers of systems. **B.** The proportion of P4 and P2 elements with defence systems deemed complete (orange, as detected by DefenseFinder or PADLOC) or putative (light orange, detected by DefenseFinder). **C.** The pangenome (black) of P4-like satellites (top) and P2-like phages (bottom) with the relative contributions of the core (blue for P4, red for P2) and the defence-associated genes (orange). **D.** Fraction of systems (detected by DefenseFinder) of the host encoded in its P4 and P2 elements. The proportions of complete defence systems encoded by P4 in hosts that have at least one P4 (leftmost panel), encoded by P2 in hosts that have at least one P2 (middle panel), or encoded either by P4 or P2 in hosts that have at least one of either element (rightmost panel) are shown in the x-axis. The size of the circles corresponds to the frequency of each (x,y) pair. **E.** Distribution of defence systems in hotspots of P4-like satellites (top) and P2-like prophages (bottom). Hotspots are defined as locations among successive core gene families and are indicated by the edges in the graphs (% indicates the proportion of defence systems at these locations). When the core genes are not successive, it reflects cases where either one of the core genes is missing, or the cores genes are arranged differently from the prototypical organization of their respective element. Colour codes in genetic loci and graphs coincide; orange indicating complete and light orange indicating putative defence systems. Grey indicates genes that are neither core nor defence-associated.

Even if this is by far the largest known dataset of P4 and P2, the pangenome saturation curves show that we are far from having uncovered the full genetic diversity of these elements (Fig 1C). Known defence genes correspond to a sizeable fraction of these pangenomes (orange lines): 34% for P4 and 17% for P2. P4 and P2 account for 11-16% of the genetic diversity of the defence systems in their hosts’ species (Fig S3). Moreover, up to 50% of the total defence systems of some genomes are encoded in their P4 or P2 elements (Fig 1D, Fig S4). This suggests that P4 and P2 are important contributors to the defence repertoire of bacteria.

Most defence systems in P4 are in a hotspot between the integrase and Psu-Delta-Sid, whilst many of the defence systems in P2 are in a hotspot between the primase (GpA) and the portal (GpQ) (Fig 1E). These are the two previously described hotspots^10^. However, other less populated defence hotspots exist between the primase and one of the *att* sites of P4 (7% of the systems in P4), between the integrase and the *att* site of P2 (3% of the systems in P2), between the integrase and the primase in P2 (8% of the systems in P2) and within the late genes of P2. Although it is atypical for integrated elements to mobilize genes between *att* sites and the integrase, this was previously observed for defence systems of PICI^21^. An additional P2 defence hotspot was found downstream of GpL (16% of the systems in P2), consistent with previous descriptions of a hotspot of accessory genes within the tail genes in the classical P2 phage^22^. Moreover, some of these P2 defence hotspots correspond to known integration sites of horizontally transferred genes in P2^23^, and to the hotspots of accessory genes described recently in more than 1000 P2-like phages^24^. In either P4 or P2, some defence systems were consistently found in a single hotspot, whilst others were found across genomic locations of the elements (Fig S5A). Of the P2 with complete defence systems, 10% have multiple defence systems across different hotspots (Fig S2C, see also genomes in Fig 1E for examples). This fraction is much lower for P4 (1%), suggesting that the accumulation of multiple defence systems occurs more often in the larger genomes of P2. These results confirm that whilst defence systems are very abundant and diverse in P4 and P2, representing a large fraction of their accessory genes, they are encoded in a small number of genomic locations.

### Rapid turnover of defence systems

We then enquired on the turnover of the defence systems, i.e. how frequently closely related elements differ in the systems they encode. The phylogenies of P4 and P2 elements, based on their core genes (see Methods), show highly similar elements often carrying unrelated defences (Fig 2A and D). This can best be appreciated by focusing on the parts of the trees with many very similar elements and inspecting the diversity of systems they contain. To quantify this effect, we compared the gene repertoire relatedness (with wGRR, see Methods) of core genes and of defence genes between pairs of elements with defence systems. Consistent with the phylogenies, we observed an overwhelmingly large proportion of pairs of the same type of elements with non-homologous defence systems (99% in P4 and 99% in P2 pairs have wGRR defence<0.05). Since this includes pairs of very similar elements (i.e., with highly similar core genes) (Fig 2B and E, right panels), it shows that defence genes change very rapidly in both P4 and P2. We observed only a few pairs of dissimilar elements with similar defence systems suggesting that convergent acquisition of defences is rare (Fig 2B and E, left panels). These patterns remain qualitatively unchanged when comparing full genomes instead of core genes and when including the putative defence systems (Fig S6). If defence systems evolved in linkage with the core genomes of P4 and P2, we would expect a linear relationship between core and defence similarity, with deviations corresponding to potential events of horizontal transfer of antiviral genes. Instead, the association between the evolution of defence systems and the core genes is almost null.

**Figure 2.**
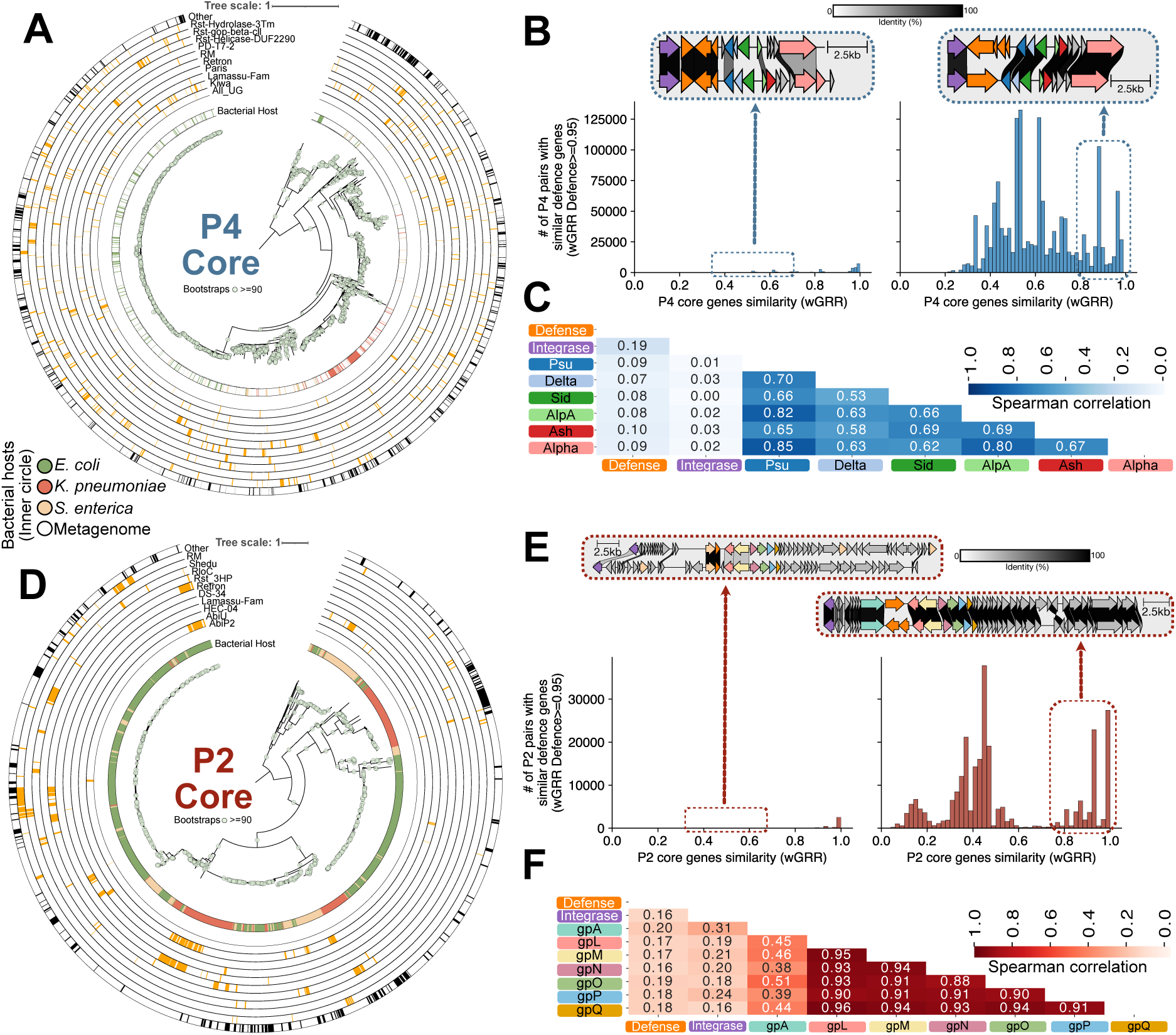
Relationship between antiviral repertoires and the core genomes of P4 and P2. **A.** Phylogeny of core genes of P4. Bootstraps of at least 90 are indicated with a circle (the larger the circle, the higher the bootstrap value). Colours in the innermost circle indicate the bacterial host of each P4 (white spaces indicate P4 from metagenomes, without an assigned host). Note that since these are dereplicated P4 genomes, some P4 might be representative of a group of elements with multiple bacterial hosts (we consider only the host of the representative genome). The ten outward circles represent the presence (orange) or absence (white) of the ten most frequent defence systems in P4 (complete defence systems, from DefenseFinder). The outermost circle indicates whether each P4 element has any other complete defence system, apart from the ten most frequent in the family of elements (black shows cases where another defence system is present). **B.** Left, number of pairs of P4 genomes with similar defence genes (wGRR≥0.95) in function of the similarity between core genes (x-axis). Right, number of pairs of P4 genomes with distinct defence genes (wGRR≤0.05) in function of the similarity between core genes (x-axis). **C.** Spearman correlation values between the wGRR of defence genes and the bitscore values from BlastP of individual core genes of P4-like satellites, for the set of all pairs of P4s. Colours of a stronger shade indicate higher Spearman correlation values. **E.** Same as in A, but for P2 genomes and their defence systems. **F.** Same as in **B**, but for P2 genomes. **G.** Same as in **C**, but for the defence and core genes of P2-like phages.

We then quantified the genetic linkage between defence genes and each individual core gene (Fig 1E). The Spearman correlations between the wGRR of defence genes and the sequence similarity of each core gene in pairs of P4 (Fig 2C and Fig S7A) and P2 (Fig 2F and Fig S7B) were all very low (≤0.2), in contrast to the correlations between core genes themselves. In summary, similar elements rarely encode similar defence genes and the latter show little genetic linkage with core genes. Both indicate very rapid turnover of defence systems in the genomes of P4 and P2.

### Pseudogenes of defence systems are rare in P4 and P2

Turnover of defence systems could occur in two ways. They could be first inactivated by mutations or indels and subsequently replaced (Deletion-reacquisition model). Alternatively, functional systems could be swapped by others before becoming defective (Swap model). The first scenario suggests that systems are lost because they are not selected for anymore, i.e., they are nearly neutral or even deleterious (costs outweigh gains). The second scenario only requires that the new defence system is at least as adaptive as the old one in the current population context. These models have different testable predictions. The deletion-reacquisition model predicts the frequent presence of pseudogenes of defence systems whereas the swap model predicts the opposite. We assessed the fraction, type, and past function of pseudogenes in P4 and P2 using PseudoFinder^25^ (see Methods). The vast majority of P4 and P2 have at least one pseudogene (94% and 98%, respectively, Fig 3), showing that defective genes remain long enough to be identifiable in large numbers in these genomes. We note that, in some cases, several identified pseudogenes can originate from a single gene. Although pseudogenes are frequently found in defence-associated loci (Fig S8), only a very small percentage of them had sequence similarity with defence-associated genes (less than 1% in P4 and c.a. 2% in P2). Many more had sequence similarity to core genes (24% in P4, typically Ash pseudogenes) and especially to unknown function genes (Fig 3 insets, see File S1 for PFAM annotations). These results suggest that defence systems are frequently swapped before being inactivated. Moreover, most pseudogenes in P4 and P2 are detected as very small intergenic sequences (c.a. 90% in either P4 or P2, Fig S9), which confirms that the putative defence systems detected in these elements are intact, likely functional genes.

**Figure 3.**
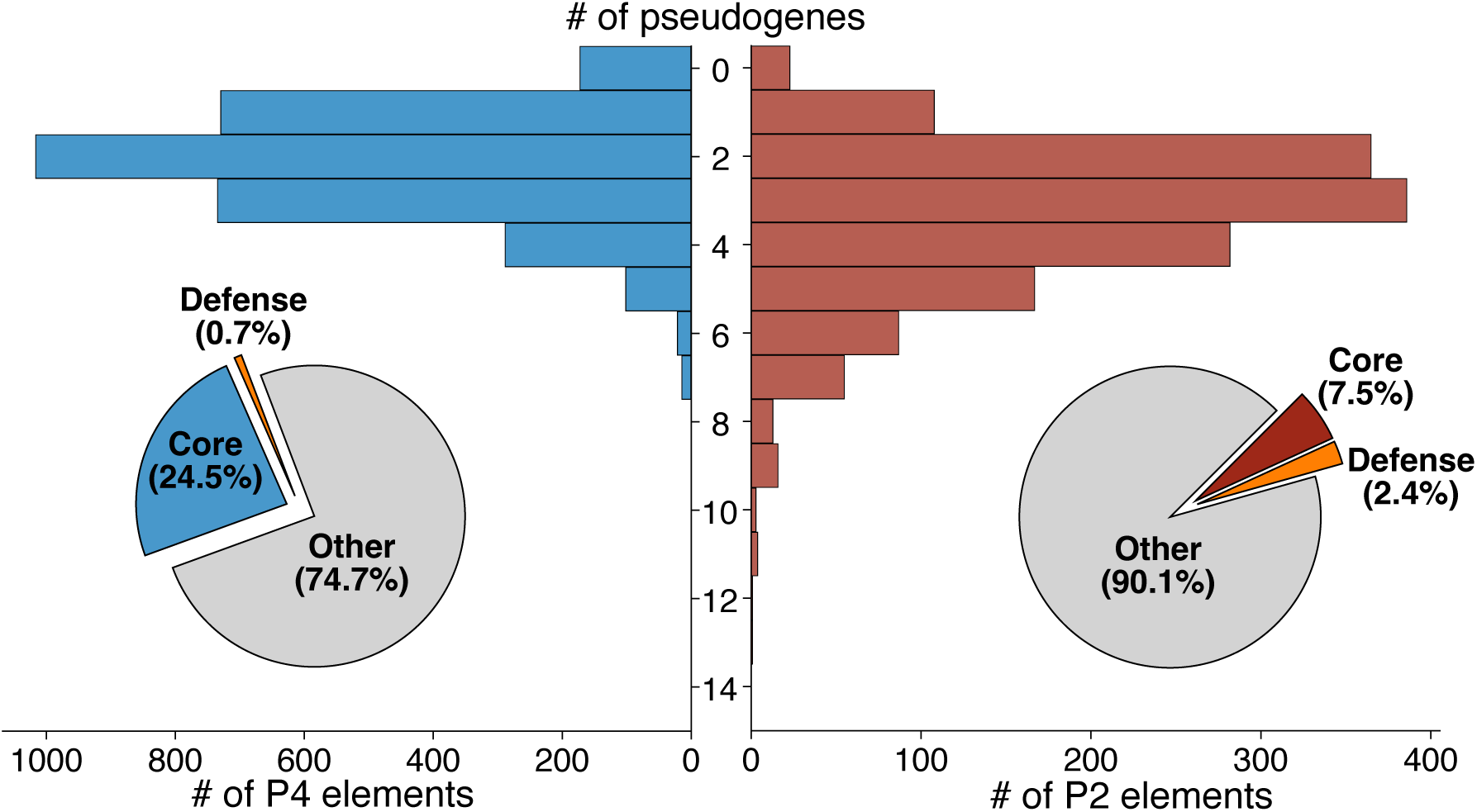
Quantification and characterization of pseudogenes in P4-like satellites and P2-like prophages. Histograms show the distribution of pseudogenes detected by PseudoFinder per P4-like satellite (left) and P2-like prophage (right). The charts as insets show the proportion of the pseudogenes annotated as defence (orange), core of each element (blue and red for P4 and P2, respectively), or with other (or no) annotations (grey).

### P4 hotspots contain numerous novel or chimeric defence systems

We found many genes of defence systems that lacked the typical cognate components in the defence hotspots (Fig S5B). These could be inactivated systems, but our observation that pseudogenes of defence systems are rare make this hypothesis unlikely. Instead, they might be functional novel systems resulting from re-assembly of others^26,27^. To test this hypothesis, we selected 11 candidate systems based on their different characteristics (Fig 4, see File S2 for a description of the putative systems): some are potential variants of known systems (e.g., a Lamassu system missing a known effector); others are mixtures of genes from different systems (e.g., two genes from the PD Lambda 2 system, and another from the Rst_gop_cll system); and others are potential minimizations of known systems (e.g., a Thoeris system with a single ThsA gene). We synthesized and cloned each putative system in a low copy plasmid under the control of a pTet promoter, which were introduced into *E. coli* MG1655. An empty plasmid was transformed to the same background as a control. We then challenged these strains with a collection of 28 virulent phages to test the antiviral protection of each putative defence system (see Methods, Fig S10, File S2). 8 out of 11 provided protection against several virulent phages (Fig 4, top 11 rows). Failure to observe protection can arise from multiple causes. The most likely explanations are: (i) the defence system is functional but does not target any of the 28 phages included in the present panel; or (ii) the system is functional in its native host but is not properly expressed, assembled, or active in the heterologous *E. coli* host used here. Interestingly, a system similar to the Lamassu variant, which did not provide protection against the phages tested here, was recently shown to be provide an anti-phage function^28^. The high success rate we obtained in detecting resistance against this limited phage panel strongly suggests that most of the putative systems not recognized as complete by either DefenseFinder or PADLOC are functional.

**Figure 4.**
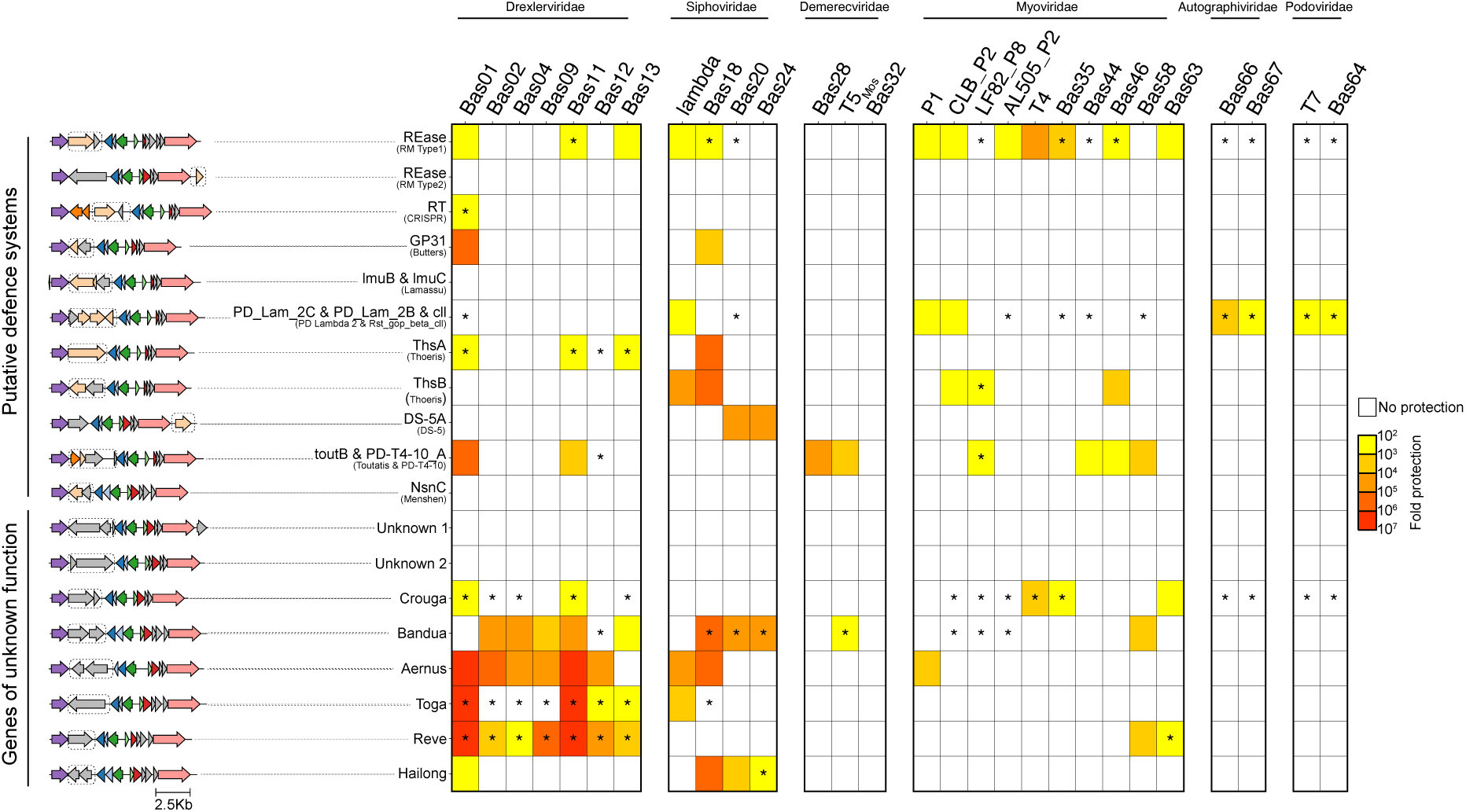
Putative and unknown defence systems in P4 provide protection against phage infection. On the left are the genomes of P4 with putative defence systems (light orange genes, top 11 rows) and the genomes of P4 with genes of unknown function in their defence loci (grey genes, bottom 8 rows). For the former, the names of the genes, as well as the complete systems with which they are associated, are shown in front of each genome. The regions synthetized in plasmids are boxed (exact sequences in Datasets E1 and E2). Heatmap of the phage resistance phenotype shows the mean fold resistance of two independent replicates of each putative and unknown system against a panel of 28 *E. coli* phages. The degree of protection is indicated by the colour (log) scale, with darker red colours indicating higher levels of protection, relative to the same bacterial background with an empty plasmid. Only the infections where both replicates show increased protection, and the mean of the two replicates is ≥ 10-fold difference, are shown as coloured squares. Asterisks indicate smaller plaque morphologies, relative to the infection of the strain with the empty plasmid. Plaque-forming units were measured for each phage on cells harbouring either a control plasmid or a defence system (Fig S10). Fold-resistance was calculated as the ratio between these two values. All systems were expressed from pTet promoter in the presence of anhydrotetracycline (aTc, 0.5 µg/ml) and measured at 37°C, except for T7-like phages, which were measured at room temperature. The raw data for the ratios system/control are shown in File S4.

Some P4 (27%) and P2 (19%) lack identifiable defence genes (Fig 1B) but have genes of unknown function at their main defence loci. We hypothesised that some of these could be novel antiviral systems. We selected at random 8 P4 satellites with genes of unknown function to test their anti-phage phenotypes, as above (Fig 4, bottom 8 rows, Fig S10). 6 out the 8 conferred protection against multiple virulent phages. phages. As previously, this high success rate indicates that most of these unannotated loci are likely *bona fide*, functional defence systems. We named the novel functional systems according to deities from Lusitanian mythology, while waiting for a uniform standardised system of nomenclature, and they are described in detail in File S3.

When possible, we characterized the novel systems using available data, sensitive sequence similarity detection programs (wjth HHPred), and structural similarity (with ESMFold and FoldSeek, File S3). The Crouga system has some evidence of being an RM-like system, with one of its proteins containing PFAM domains (ResIII, DEAD and Helicase C) associated with other RM systems^29^. The Bandua system has two proteins with domains typically associated with immune functions (a nuclease and a kinase, respectively). Neither protein is recognized by current antiviral annotation pipelines. The Aernus system has one protein annotated with kinase and nuclease domains, and a second one annotated as a mannitol repressor. It is unknown at this point if the latter has any impact in antiviral capabilities. The single protein in the Toga system has several hits with protein domains associated (through FoldSeek) with the Gao_Ppl^30^ and the SMC defence systems^31^, albeit not being recognized by any of the current models for either system. The Reve system is another single gene system, with ATPase and KAP family P-loop domains^32^. Finally, the last system we tested has two proteins, one with pentapeptide repeats and an IRK potassium channel domain, which could be involved in membrane depolarization, while the other protein contains a Nucleotidyltransferase like domain and a HEPN-like domain, suggesting it acts on host or phage RNA. A system with similar components was described by others during the course of this study as Hailong^33^, which is currently not detected by DefenseFinder or PADLOC. Hence, and despite being part of the non-defence annotated genes in P4, we refer to it as Hailong to avoid introducing redundant nomenclature.

The novel systems described above, together with the unannotated Hailong system, are clear evidence that at least some genes of unknown function in P4 have antiviral phenotypes, with most (e.g., Bandua, Aernus, Toga and Reve) providing strong protection against Drexlerviridae and/or Siphoviridae phages). We thus wondered whether other genes in the defence-less elements could have similar properties. For this, we collected all the genes of unknown function from P2 and P4 detected only in complete genomes, to use the information on their larger chromosomal context, and analysed them using a recent machine learning approach to detect antiviral domains (see Methods). We observed a significant signal for the association of antiviral defences in 30% of the proteins of unknown function in P4 (62%, with the method that considers the genomic context), and in 15% of the unknown proteins in P2 (32%, with the method that considers the genomic context) (Fig S11). We note that all the confirmed novel systems are part of those predicted to have an antiviral function, due to their genetic context. The computational predictions, together with experimental validations, demonstrate that P4-like satellites constitute major loci for both established and yet-to-be-discovered functional defence systems. Moreover, many of the proteins of unknown function that cluster in P4- and P2-associated defence hotspots are predicted to encode antiviral activities.

### P4 and P2 exchange defences with other MGEs but not with each other

If defence systems are constantly swapped in both P4 and P2, where do these genes come from? To address this question, we searched for closely related homologs (>85% identity, see Methods) of defence genes from either P4 or P2 in 32798 bacterial and in 27022 phage genomes. For 6% of defence genes in P4, and for 2% of the defence genes in P2, we found at most one homolog, indicating that some of these defence genes are likely variants specific to the P4 and P2 in our dataset (Fig S12). Many of the remaining P4 and P2 defence genes have homologs in multiple genomic contexts (Fig 5A). Similar results were found when including the putative novel defence systems from the first section (File S5, Fig S13). To avoid comparing elements that might have similar systems because of recent common ancestry, we excluded the pairs of very similar (wGRR>=0.95) P4 or P2 elements. Nevertheless, the analysis reveals that most homologs of defence genes of P4 (76%) and P2 (47%) are in bacterial regions predicted to be respectively P4-like satellites and P2 phages. These include P4 and P2 in bacterial hosts (e.g., *Citrobacter*, *Enterobacter* or *Pectobacterium*, Fig S14) where these elements’ delimitation in bacterial chromosomes was not possible for lack of information on their *att* sites (called P2* and P4*, see Methods). These elements had hence not been included in our focal dataset of phages and satellites. P2 and P4 often co-occur in genomes and are induced and replicated at the same time. One could thus expect that they would exchange defence systems. Surprisingly, we found only one case of closely related defence genes between P4 and P2, even when including all putative defence systems (Fig S15). Hence, hitchers and helpers have not recently exchanged defence systems.

**Figure 5.**
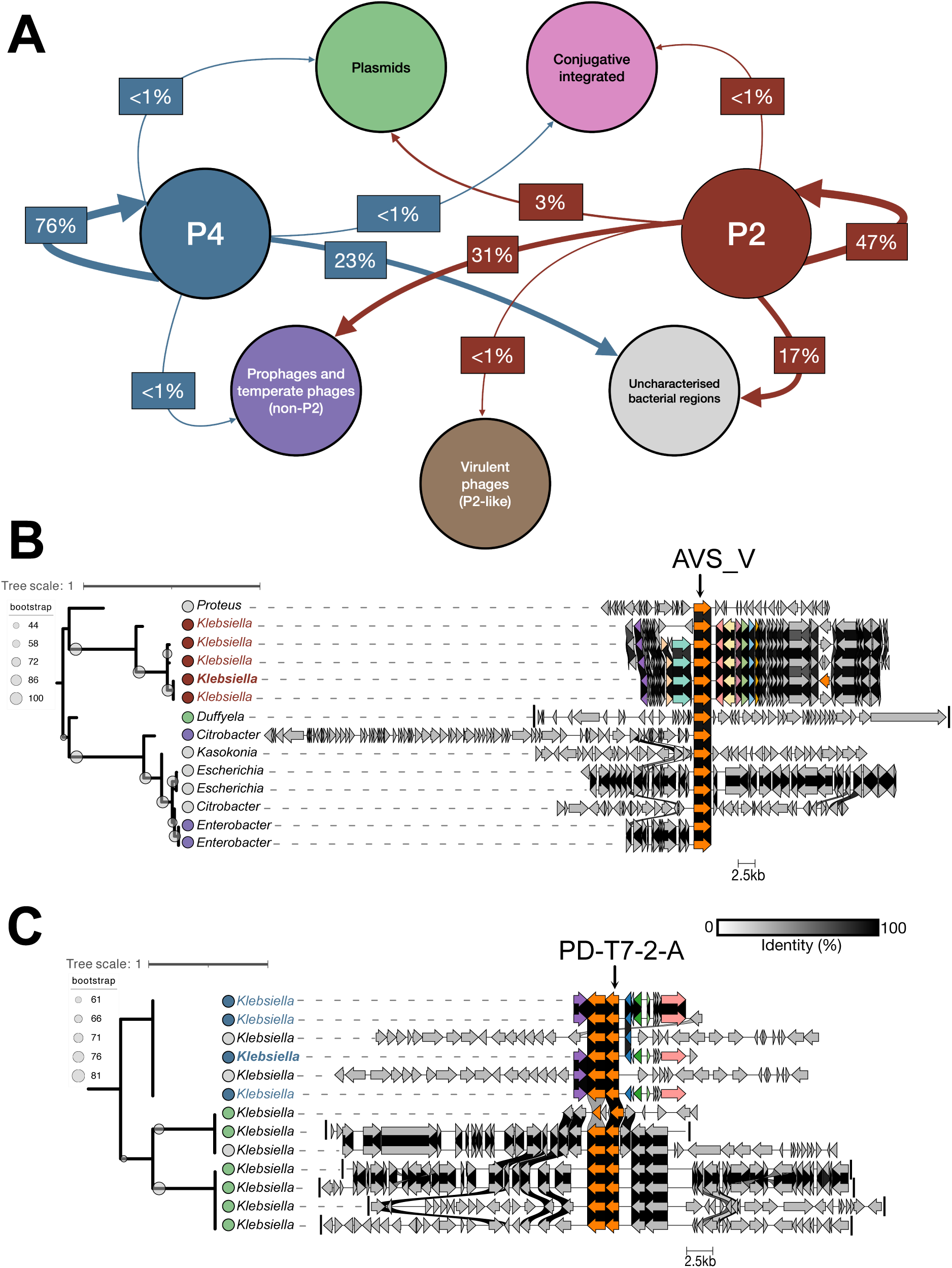
Highly similar homologs of defence genes of P4 and P2 are found in completely different genomic contexts. **A.** Graph based representation of the number of instances where homologs of defence genes (from complete defence systems) in P4 and P2 are found in different genomic contexts. These include other MGEs (plasmids, phage-plasmids, conjugative integrated elements, prophages and virulent phages) or regions of the bacterial genomes that are uncharacteristic. Thickness of the arrows represents the proportion of homologous matches detected between defence genes from P4 or P2 and the other different genomic contexts (thicker arrows correspond to a higher proportion of matches). The exact values for each combination are shown in File S5. **B.** Phylogeny of the AVS_V gene, comprised of the gene found as a defence system in a focal P2 in *Klebsiella*, and its homologs detected in either other P2s (red circles at the tree leaves), plasmids (green circles), prophages (violet circles) or uncharacteristic genomic regions (grey circles). The genes corresponding to each of the elements (or to the segments that include the uncharacteristic genomic regions as grey circles) are shown in front of the tree, with larger plasmids truncated for visibility. **C.** Phylogeny of the PD-T7-2-A gene, comprised of the gene found in a PD-T7-2 defence system in a focal P4 in *Klebsiella*, and its homologs detected in either other P4s (blue circles at the tree leaves), plasmids or uncharacteristic genomic regions. The genes corresponding to each of the elements (or to the segments that include the uncharacteristic genomic regions) are shown in front of the tree, with larger plasmids truncated for visibility.

The highly similar homologs of P4 and P2 defence genes found in bacterial regions that are not identified as P4 or P2-like could be the result of recent genetic exchanges with other MGEs. We found no homologs in the other families of satellites known in Enterobacteria (PICI and cfPICI^9,17^,File S5). Both P4 and P2 defence homologs were found in (non-P2 like) prophages, albeit this was much more frequent for P2, which also has defence homologs in 31 temperate and 11 virulent phage genomes (all P2-like). Chromosomal regions with integrative conjugative elements^34^, which are known to encode antiviral systems^35^, account for very few of the homologs of P4 or P2 defence genes. P4 and especially P2 defence systems have relatively more homologs in plasmids. Some of the latter could correspond to phage-plasmids (P-Ps), since they are plasmid replicons identified as phages by geNomad^36^, although they are not part of the current dataset of known P-Ps^37^. The MGEs and bacterial regions with homologs of defence genes of P4 and P2 are sometimes within bacteria that are phylogenetically distant from the hosts of P4 and P2, such as *Pseudomonas* or *Vibrio* (Fig S14). These results suggest that exchange of defence systems tends to occur between similar types of elements, but can also involve distinct MGEs.

Genetic exchanges with other MGEs may allow to introduce novel types of defences in P2 or P4, which can then be exchanged within each group of elements. For instance, the phylogeny of a variant of the AVS_V system, detected in P2 from *K. pneumoniae*, includes homologs in non-P2 prophages and uncharacteristic genomic regions from different bacterial hosts, that form a separate clade (Fig 5B). Intermingled with the two main clades are two terminal branches, corresponding to an uncharacteristic genomic region in *Proteus* and a single plasmid in *Du<yella*. The phylogeny suggests the acquisition of the system by P2 was recent. It was then followed by the dissemination of this variant of AVS_V within *K. pneumoniae* in P2 phages. Another defence system detected in P2, Lit, is mostly found in plasmids and uncharacteristic genomic regions. While being rare in prophages, a homolog is found in a single P2-like prophage (Fig S16A). Given its singularity, and the scarcity of this type of systems in P2 (Fig 1A), this likely illustrates a recent acquisition of the Lit system by this P2.

Exchange of defence systems between different types of MGEs also broadens the mechanisms of dissemination of the former within bacterial species. For instance, the phylogeny of a variant of the PD-T7-2 A is consistent with a unique, recent exchange of this system between P4 and plasmids (whose direction cannot be ascertained from this data alone). The system in the plasmids has diversified in two separate genetic variants that disseminated in *K. pneumoniae* (Fig 5C). The phylogeny of the other gene of the system (PD-T7-2 B) shows yet another variant in P4 that is disseminating the defence system in *Escherichia* and *Klebsiella* hosts, with a transfer to plasmids in the latter (Fig S16B). Hence, events of exchange between MGEs show the establishment and further diversification of defence homologs in different MGEs, allowing the spread of defence systems within and across species through different mechanisms of horizontal gene transfer.

### Cryptic genomic islands have homologs of defence genes from P4 and P2

A large fraction of homologs of P4s or P2s identifiable defence systems are found in bacterial <u>u</u>ncharacteristic <u>g</u>enomic regions (UGRs) that do not correspond to known MGEs. Their genomic context (20 genes each side) often contained tRNAs genes, which are well-known points of integration of MGEs^38^ (Fig S17A). Functional annotation with EggNOG Mapper and PFAM revealed that some of these genes are typical from P4 or prophages (Fig S17C and D, see blue and red dots). These regions also contained many pseudogenes of proteins typical of MGEs (Fig S17A and B). Since these results suggested that some UGRs could be defective satellites or phages, we systematically annotated them using the core markers of P4 and P2 (see Methods) and PseudoFinder. In total, 745 URGs encode highly similar homologs of bacterial antiviral genes detected in P4s (Fig 6A, File S5) and many of them (27%) have at least one P4-specific marker(s). All 745 UGRs have at least one pseudogene, and about half have pseudogenes (typically intergenic) homologous to a P4 core gene. Conversely, in the 775 UGRs that encode highly similar homologs of bacterial defence genes detected in P2s, P2-like markers are found in ca. 20% of them. Although almost all (>98%) of these UGRs have at least one pseudogene, only 5% of them correspond to a P2-like core gene. The above observations remain qualitatively similar when including the putative defence systems (Fig S18). UGRs with P4 or P2 core markers are thus very likely defective versions of these elements. Within them, we found numerous pseudogenes of mobility functions, but the neighbouring defence genes remained almost identical to those of intact elements, and are thus likely functional (Fig 6B-C). These results mean that defective P4 and P2 either engaged in genetic exchanges of defence systems with their functional counterparts, or are non-mobile satellites and prophages with stabilized (and conserved) defence genes. Moreover, and consistent with the absence of genetic exchanges between hitcher and helper (Fig 5), homologs of defence genes in P4 and P2 are also not found in defective elements of the opposite type.

**Figure 6.**
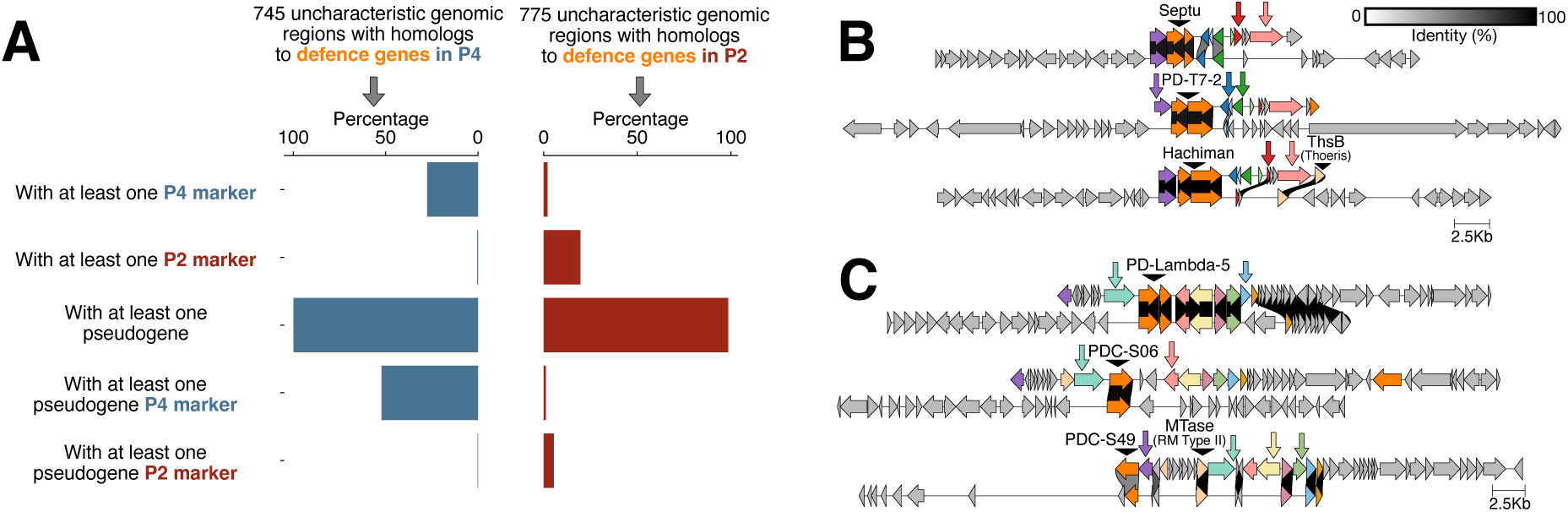
Quantification of pseudogenes and P4-like/P2-like markers in uncharacteristic genomic regions with homologs to defence genes encoded by P4 and P2. **A.** Characterization of uncharacteristic genomic regions (URGs) that encode homologs to defence genes found in P4 (left bars, blue) and URGs that encode homologs to defence genes found in P2 (right bars, red). We quantified the proportion of these URGs that encode P4/P2 markers predicted as pseudogenes by PseudoFinder (last two bars). **B.** Examples of P4-like satellites, and the homology of their defence systems with URGs that contain incomplete (likely defective) P4-like satellites. In each genomic comparison, the coloured arrows indicate P4-like core genes that were detected as intergenic pseudogenes in the URGs. **C.** Examples of P2-like prophages, and the homology of their defence systems with URGs that contain incomplete (likely defective) P2-like prophages. In each genomic comparison, the coloured arrows indicate P2-like core genes that were detected as intergenic pseudogenes in the genomic regions.

Almost 50% of UGRs with homologs to P4 defences and 80% of the UGRs with homologs to P2 defences lack any P4- and P2-specific markers, considering both the genes and pseudogenes within these regions. They are thus cryptic genomic islands, completely distinct from P4/P2, and of unknown mobility. The detection of defence homologs in such dissimilar islands is consistent with the observed putative exchanges of P4 and P2 defences with MGEs of different types, where we do not expect to find large regions of homology (Fig 5).

## DISCUSSION

The systematic detection of defence genes in prophages and satellites is a strong statement on the selective importance of antiviral functions for the ecology and evolution of these MGEs. Given the prevalence of P4 and P2 in Enterobacteria^17^, and the large proportion of their accessory genes that we show here to be dedicated to defence, these MGEs constitute a significant fraction the bacterial repertoire of defences. Yet, the elements’ pangenome dynamics show that the sampling of their natural diversity is not yet exhausted, even when considering, for the first time, the vast diversity of P4 elements in metagenomes using Logan^18^. Our experimental validation of anti-phage activity for 13 novel systems encoded by P4 confirms that the contribution of defence functions to the pangenomes of P4 and P2, as well as to the antiviral repertoire of their bacterial hosts, will keep surprising us.

The diversity of antiviral functions in P4 and P2 is driven by events of genetic swaps at defence-specific hotspots, enlightening the underlying processes of natural selection in MGE-defence associations. A novel system will fix in a population of MGEs if it has adaptive value. Targeted phages may, over time, adapt to resist this novel system, reducing the selective advantage of retaining it in the MGE genome. It has remained unclear if this process typically runs until the defence systems become neutral (or even deleterious, e.g., due to autoimmunity^39^) or if defences are replaced beforehand. Our observation of rapid replacement of the systems and the lack of defence-associated pseudogenes at these loci suggest that recombination quickly swaps functional systems without the latter passing by a period of inactivation. Although many small intergenic regions predicted as pseudogenes are in defence loci, they are likely remnants of an intense process of recombination. What little may remain of antiviral genes in these regions lost any recognisable homology with known defences, even if they could potentially give rise to novel defence genes^40^. These findings have two implications. First, defence systems may be lost when they are still adaptive: it suffices that the novel system is more adaptive for the swap to be fixed in the population by natural selection. Second, it suggests that systems spend little time, if any, as neutral components of the MGE genome.

P4 and P2 exchange their defence systems with elements of the same type and, less frequently, with other MGEs. Yet, they do not exchange with each other. This is puzzling because P4 and P2 frequently co-occur within bacterial hosts, and are co-induced, replicate and package their DNA simultaneously^41^. This should provide plenty of opportunities for genetic exchanges^9,42^, particularly since phage induction, replication and recombination are mechanistically concomitant^43,44^. Three factors may explain this result. First, exchanges by homologous recombination require high sequence identity that is lacking between P2 and P4^45^. Yet, we do observe exchanges between these elements and other unrelated MGEs, like plasmids. Second, because P4 and P2 often co-occur in genomes, there might be selection against the presence of redundant defences in the two elements. Different types of defences in P2 and P4 would maximize the range of anti-phage defence of the consortium. Identical defences would be redundant, and one set could be easily lost by genetic drift. Third, if P2 have defence systems to prevent P4 infection, or to avoid the hijacking of P2 particles by P4, these systems might be counter-selected in P4. Conversely, P4 might carry defence systems that increase the fitness of P4 by decreasing the ability of P2 to use its own capsids. If they exist, such genes are not expected to be acquired by P2 phages.

Despite the lack of exchanges between P2 and P4, we found very similar defence systems in distinct MGEs or genomic islands, suggesting that genetic exchanges between very different elements is possible. It is difficult to estimate the direction of these exchanges, although in some cases the defence was very likely acquired by P4 or P2 from other MGEs. When a phage or a satellite acquires a defence gene from a plasmid, the system can now be exchanged more easily with other satellites or phages^46^. These exchanges can also contribute to the flow of antiviral genes between phylogenetically distant bacteria, if the MGEs involved in these exchanges have different host ranges. We also found defence homologs in defective MGEs. It will be interesting to investigate if this corresponds to their domestication, resulting in the stabilization of the defence genes in bacterial genomes, or if defective elements still actively swap their defences with other MGEs.

Defence genes in P4 and P2 have evolutionary histories that are strikingly different from the rest of the elements’ genomes. Antiviral defences, in accordance with the idea that they are “guns for hire”, change genomic contexts of MGEs as quasi-independent entities^47^. Yet, they do so in ways that resemble the change of bacterial chromosome contexts by MGEs: frequent recombination breaks genetic linkage between defence genes and the core genes of the elements, like MGEs are unlinked from the bacterial chromosome lineage^48^; defence genes integrate predictable (hotspot) locations, like MGEs in bacterial chromosomes^49^ (through a presumably different mechanism); the modularity of defence systems can promote the emergence of hybrids through recombination, resembling the modularity of the evolution of prophages^50,51^; defence genes are a large fraction of the satellite and the phage’s pangenomes, similar to the contribution of MGEs to bacterial pangenomes^52^; and defence genes provide accessory functions to both phages and satellites, which in turn do the same to their bacterial hosts^2^. The key difference is that defence systems are not known to encode mobilization functions. Instead, our results suggest that P2 and P4 hotspots function as factories of novel systems by recombination processes that still need to be elucidated. It also remains to clarify how mutations and natural selection act on these chimeras to eventually lead to their fixation as functional hybrid systems. In summary, our work shows that P2 and P4 can be seen as genetic platforms that accumulate, diversify, and then disseminate bacterial defence systems across microbial populations.

## MATERIALS AND METHODS

### Genomic and metagenomic datasets

We retrieved all the complete genomes of the NCBI non-redundant RefSeq database (ftp://ftp.ncbi.nlm.nih.gov/genomes/refseq/, last accessed in May 2023), which consists of 32799 complete bacterial genomes. The metagenomes used for the detection of P4-elements were retrieved by aligning a query set of 538 clustered P4 protein sequences to Logan v1.1 contigs using DIAMOND2^53^, as described in the Logan manuscript^18^. We also retrieved from GenBank a dataset with 27022 complete bacteriophage genomes (retrieved March 2024). The lifestyle of each of these bacteriophages was inferred using Bacphlip^54^.

### Identification and delimitation of genomic regions of P4-like satellites or P2-like prophages in complete bacterial genomes

P4-like satellites were initially detected from the RefSeq bacterial genomes as described previously^17^. This resulted in 3437 P4-like genomes of Type A or B. All prophages encoded by bacterial genomes were detected using geNomad (version 1.5.2, default parameters)^36^. In total, we detected 88288 prophages in the RefSeq bacterial genomes. We then designed a MacSyFinder^55^ model corresponding to a prototypical P2-like prophage. The components of the model include the GpQ-L operon, which is conserved in P2-like prophages^56^, as well as an integrase and a primase (GpA). This model was directly applied to the prophages detected by geNomad, resulting in 8165 P2-like prophages of Type A, B or C. The latter two types of P2-like prophages are often missing either the integrase, or the primase. We speculate that these two components are either less conserved than the profiles used for the GpQ-L operon, or excluded by geNomad during the predicted prophage region. The proteomes of these elements were retrieved by extracting all the genes within the farthest core components. This general set of non-att delimited 3437 P4-like satellites and 8165 P2-like prophages constitutes what we name, respectively, the broad P4* and P2* categories.

A stricter definition of the genomic regions corresponding to P4 and P2 was achieved by identifying *att* sites (attL and attR) at the borders of these elements in three bacterial species, *Escherichia coli* (Ec), *Klebsiella pneumoniae* (Kp), and *Salmonella enterica* (Se), which constitute the most abundant hosts of P4 and P2 elements^17^. To do so, we started by identifying genomic spots containing P2 or P4 core genes across bacterial chromosomes. The genomic positions of P4 and P2 elements described above were mapped relative to persistent genes in the three host species (Ec: 2527 genomes, Kp: 1510 genomes, and Se:1256 genomes) to facilitate MGE delimitation and comparison across genomes. The pangenome of each species (i.e., the full bacterial species gene repertoire, not to be confused with the P2/P4 pangenome) was computed using the pangenome module of PanACoTA v1.4.1-dev^57^, applying a minimum sequence identity and coverage threshold of 80%. Persistent gene families—those present in a single copy in at least 90% of genomes—were extracted using the corepers module of PanACoTA. Intervals were defined as the regions between two consecutive persistent genes within genomes. P2 and P4 core genes are not persistent across species and are confined to these intervals, providing an upper bound for their genomic positions. In total, 6,008 intervals containing P2 or P4 core genes (excluding integrases) were identified across the three species (2258 in *Ec*, 2,66 in *Kp*, and 1384 in *Se*). Intervals from genomes of the same species were grouped into genomic spots if flanked by the same pair of persistent gene families, allowing comparison of equivalent regions across genomes. The number of genomic spots occupied by P2 and P4 was generally low in each species (30 in *Ec*, 16 in *Kp*, and 21 in *Se*).

P2 and P4 integrate the bacterial chromosome using a Tyrosine recombinase (integrase), which mediates site-specific recombination between the host *attB* site and the MGE *attP* site. Integration duplicates part of *attB* at the element edges, generating *attL* and *attR*^58^, which can be identified by sequence similarity within genomic spots. For each spot described above, the shortest interval observed in the species was extracted as the ancestral MGE-free state containing *attB*; when this assumption does not hold, *att* sites cannot be reliably identified. Each such interval was then compared using sequence alignments with longer intervals containing P2 or P4 elements, expected to harbour short repeats of *attB* at their edges. To ensure at least two significant sequence alignments per comparison and to account for possible integrations within persistent genes, intervals were extracted including the flanking persistent genes and oriented so that these genes point in the same direction, restricting the search to the positive strand for computational efficiency. Sequence similarity between the shortest interval (query, *attB*) and each longer interval (subject, candidate *attL/attR*) within the same spot was screened using BLASTn of BLAST 2.9.0+^59^ (-evalue 0.001 -strand plus). Alignments with ≥80% identity were retained, and redundant hits fully contained within longer alignments were discarded. Pairs of hits on the subject aligning to overlapping regions of the query were interpreted as duplications associated with integration. The overlapping region on the query defines the putative *attB* site, and the corresponding regions on the subject define *attL* and *attR*. Pairs of *att* sites flanking all P4/P2 core genes were then selected to precisely predict element boundaries, resulting in the delimitation of 2175 elements in *Ec*, 2230 in *Kp*, and 1357 in *Se,* corresponding to 96 % of intervals containing P2/P4 core genes. This approach does not rely on genomic context, such as the presence or position of tyrosine recombinase genes (unlike methods such as TIGER^60^). Although integrases were not considered while detecting P4 and P2 spots, they are present in nearly all delimited elements – with only 76 exceptions - and in most cases (5638), the integrase is located within 2kb of an element boundary, demonstrating the robustness of our method. Att site pairs (sequences), element size, and integrase presence are provided in File S6.

From the P4/P2 intervals with an *att* identification, we further filtered the elements according to their size (<20kb for P4 and <50kb for P2) and discarded P4 that were lacking more than two core genes (i.e, only considered P4 of Type A or B, according to SatelliteFinder). Delimiting and filtering the elements in the 3 main bacterial species, resulted in a total of 2168 P4-like satellites and 3574 P2-like prophages originating from complete bacterial genomes.

### Identification and delimitation of genomic regions of metagenomes that correspond to P4-like satellites

To detect P4-like satellites in the metagenomic contigs, the latter were initially processed as described previously in^18^. The 7 P4 core genes were detected in a set of 210895 “P4-like contigs”. An initial step of dereplication of data was performed in these full contigs, using MMSeqs2^61^ with an identity cutoff of 0.999 and a (default) minimum of 80% coverage. This resulted in 16205 unique “P4-like contigs”. The collection of *att* sites detected for the P4-like satellites in the 3 main bacterial species (described above) were then used to subsequently delimit the regions in each contig that correspond to a P4-like satellite. Briefly, we used blastn with the *blastn-short* option to detect the *att* sites in each contig, retaining matches with ≥ 70% identity. To consider possible variation in *att* sites, we did not enforce the match with both a specific *attL* and *attR* sequence, but instead searched for either one of these sequences (i.e., two *attL* or two *attR* sequences would suffice). We then extracted the shortest regions that correspond to nucleic sequences that are surrounded by two of these *att* sequences, requesting a minimum of 5kb and a maximum of 20kb in length for the whole *att*-defined region, according to the expected distribution of sizes of P4-like elements. This resulted in 6039 unique P4-like replicons. To confirm that these regions correspond to the expected P4-like satellites with enough components, we first used Prodigal to extract the proteins, and then used SatelliteFinder to characterize the putative P4s. This resulted in a total of 6026 P4-like satellites, where most sequences (6001) correspond to P4-like satellites of Type A or B (Fig S1). This suggests that our delimitation of P4-like regions using the known *att*-sites, whilst likely conservative and resulting in the discarding of some valid elements, is appropriate to properly define the regions of P4-like satellites.

### Dereplication of P4 and P2 genomes

To dereplicate the final dataset used for the analyses, we merged the nucleotide sequences from the *att*-defined P4-like satellites *in E. coli*, *K. pneumoniae* and *S. enterica* with all the *att*-defined P4-like contigs from the metagenomes. We then clustered these replicons using MMSeqs2 with the same identity cutoff as before (0.999) but with a stricter 99% coverage threshold, to avoid the merging of elements that have only one gene that is distinct. A similar approach was done for the *att*-defined P2-like prophages in the 3 main bacterial species. This resulted in a final dataset of 3085 dereplicated P4-like satellites and 1511 dereplicated P2-like prophages, which we use for all the analysis described in the manuscript.

### Identification of defence genes and systems

The detection of defence genes and systems were done with DefenseFinder version 2.0.0 (defense-finder-models database v2.02)^19^, using the default options. We included the flag -a to detect the anti-defence systems as well, albeit none were detected in P4-like satellites, and very few in P2-like prophage (see main text for details). We used PADLOC version 2.0.0^20^ to complement the detection of complete defence systems by DefenseFinder. To retrieve genes associated with putative defence systems, we extracted the best hit for each gene from the *.hmmer.tsv output file of DefenseFinder.

To identify the gene repertoire of defence systems in the bacterial hosts of P4 (restricted to P4 detected in complete bacterial genomes) and P2 elements, we identified the defence systems in the bacterial genomes of the three main species (*E. coli, K. pneumoniae* and *S. enterica*) that encoded at least one P4 and/or one P2, using the same approaches as above. The defence gene families in bacterial hosts were computed by collecting, for each species, the genes of the defence systems (either in DefenseFinder or PADLOC) and clustering all the corresponding protein sequences using MMSeqs2 (again, the clustering was performed per each of the three species, and for each dataset, DefenseFinder or PADLOC individually). A defence gene family was considered to be associated with P4 or with P2 if at least one of the genes of the family is within an *att*-defined P4 or P2 genome of the three main species.

### Calculation of P4 and P2 pangenome accumulation curves

We first clustered at 40% identity all proteins of the dereplicated P4 satellites, and all proteins of the dereplicated P2 prophages, using the MMSeq2 cluster function, with the parameters cluster_mode 1 and the default coverage. Then, for each category of elements (P4 or P2), we randomized the order of the elements and calculated the number of new gene families added by each consecutive element, annotating whether their representative sequences correspond to a defence-associated gene or a core gene of the respective element.

### Calculation of genomic similarity between P4-like satellites and between P2-like prophages

We searched for sequence similarity between all proteins of satellites using MMseqs2 with the sensitivity parameter set at 7.5 to align all versus all proteins. For specific datasets (core or defence genes), the datasets were limited to the genes that were annotated with the specific core markers or identified as defence-associated. We then performed the combinations between different datasets as well (e.g., defence vs core or defence vs whole proteome). The outputs of MMSeqs2 were converted to the blast format and we kept for analysis the hits respecting the following thresholds: e-value lower than 0.0001, at least 35% identity, and a coverage of at least 50% of the proteins (since MMSeqs2 searches for local similarity). The hits were used to retrieve the bi-directional best hits between pair of genomes, which were in turn used to compute a score of gene repertoire relatedness weighted by sequence identity:

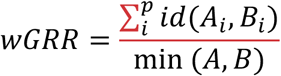

where A_i_ and B_i_ is the pair *I* of homologous proteins present in *A* and *B*, id(A_i_,B_i_) is their sequence identity in the local alignment, and min(*A*,*B*) is the number of proteins of the genome encoding the fewest proteins (*A* or *B*). wGRR is the fraction of bi-directional best hits between two genomes weighted by the sequence identity of the homologs. It varies between zero (no bi-directional best hits) and one (all genes of the smallest genome have an identical homolog in the largest genome). Hence, a wGRR=1 means that the two elements are identical (or one is entirely contained within the other), whereas wGRR=0 means the elements have no homologous protein-coding genes. wGRR integrates information on the frequency of homologs and sequence identity. For example, when the smallest genome has 10 proteins, a wGRR of 0.2 can result from two identical homologs or five homologs each with a lower sequence similarity (40%). All wGRR calculations in the manuscript were obtained through GRIS version 2 (https://gitlab.pasteur.fr/jgugliel/gris).

### Calculation of the pairwise similarity of core genes of P4 or P2

To calculate the pairwise similarity between each core gene, we first collected and concatenated in a fasta file all the individual core genes of P4-like satellites (7 core genes) and P2-like prophages (7 core genes). Each of these files with concatenated proteins was simultaneously used as both query and target database in Diamond BlastP, in order to retrieve the best pairwise matches. The parameters used were the following: --query-cover 50, --ultra-sensitive and --max-target-seqs 0. We then retrieved the best matches for each core gene between each pair of elements.

### Identification of pseudogenes

Pseudogenes were detected using PseudoFinder version 1.1.0^25^, using the default parameters. We first generated the GenBank files from the sequences to be analysed (P4-like satellites, P2-like prophages, or uncharacteristic genomic regions of bacterial genomes). We used as a reference database the complete set of bacterial proteins from the RefSeq bacterial collection used in this study (see Bacterial genomic and metagenomic datasets). We then analysed the putative pseudogenes reported by PseudoFinder either as an existing ORFs or as an intergenic region predicted to be a pseudogene. To functionally annotate the latter, we used the ORF from the RefSeq bacterial dataset that better matches each intergenic sequenced (as reported by PseudoFinder, using BlastX). The putative pseudogenes (ORFs and intergenic) were then functionally annotated with the HMM profiles corresponding to the P4 and P2 core genes and analysed with DefenseFinder and PADLOC to detect defence-associated genes. Additionally, we annotated the pseudogenes using EggNoggMapper version 2.1.12^62^, with the -m diamond parameter and with all the other parameters left as default.

### Cloning of candidate defence systems

Putative defence systems, as well as unknown P4 genes, were synthesized with their native promoter and cloned into the low-copy plasmid pFR66, which carries a pSC101 origin of replication, a kanamycin resistance cassette, and a superfolder GFP gene under the control of a pTet promoter^10^. Thus, the GFP was replaced by the candidate defence systems, placing them under the control of the anhydrotetracycline (aTc) inducible pTet promoter. Fifty µl of *E. coli* K-12 MG1655 chemically competent cells prepared by rubidium chloride method were transformed by heat shock at 42°C with one µl of plasmid carrying the candidate defence system. After 1 h of recovery at 37°C in LB (Lysogeny broth) medium, transformants were selected on LB plates supplemented with kananamycin (Kan) at 50 µg/ml. All constructions were verified by Nanopore sequencing. The fragments of candidate defence genes are shown in Dataset E1, whilst the complete plasmid sequences are shown in Dataset E2.

### Phage amplification

The panel of 28 phages tested, including both classical model phages and a representative selection of phages from the BASEL collection^63^, as well as the T5_Mos_ phage^64^, was amplified on *E. coli* K-12 MG1655. For amplification of phage stocks from - 80°C, a frozen piece of phage stock mixed with 200 µl of an overnight culture of *E. coli* and 20 mL of warm (56°C) LB + CaCl2 5 mM + 0.5% agar was poured onto square plates (12 x 12 cm) containing LB + CaCl2 5 mM + 1% agar and incubated overnight at 37°C except for the T7-like phages that were incubated at room temperature. The top-agar layer was recovered with 1 mL of PBS 1X from plates on which confluent lysis was observed and transferred into a 50 mL conical tube. The top agar was then disrupted by vortexing until broken into small pieces. Tubes were then left to incubate 10 min at room temperature before centrifugation at 4,000 g for 5 min. Finally, the supernatant was recovered and filtered with a 0.45 µM filter. The phage lysates were stored at 4°C. To determine the titer of the phage lysates, the lysates were serially diluted and spotted onto lawns of *E. coli* MG1655.

### Phage plaque assays

*E. coli* K-12 MG1655 strains carrying each putative defence system or the control plasmid pFR66 were grown overnight in LB supplemented with Kan 50 µg/ml. Bacterial lawns were prepared by mixing 200 µL of a stationary culture with 100 µl of 1 mM CaCl2 and 20 mL of LB containing 0.5% agar, and the mixture was poured onto square plates (12 x 12 cm) of LB supplemented with Kan 50 µg/ml with or without 0.5 µg/ml aTc. Serial dilutions of high-titre (>10^8^ pfu/mL) phage stocks were spotted onto each plate and incubated overnight at 37°C or at room temperature for T7-like phages. The next day, plaques were counted, and fold resistance was calculated as the number of plaques in the presence of each system divided by the number of plaques on the control plate. When plaques were too small to be counted individually, the highest-concentration dilution showing no visible plaques was conservatively scored as containing a single plaque.

### Machine learning detection of antiviral domains in P4-like satellites and annotation of novel defence systems

To annotate the candidate defence systems and characterize potential antiviral domains, sequence and structural homology analyses were performed. Sequence homology was first assessed using Pfam-A annotation using PyHMMER^65^ (v0.11), a Pyhton library binding to HMMER^66^ (v3.4). To annotate protein regions without Pfam annotations, HHpred^67^ was used with 3 iterations on Pfam-A (online version) to identify conserved domains via profile-to-profile Hidden Markov Model (HMM) comparisons. Structural predictions were used for both homology detection and representation. For homology detection, 3D protein models were generated using ESM-Fold, and the resulting PDB files were processed through the Foldseek web server^68^ to identify structural homologs within the PDB and AlphaFold databases. In the second approach, AlphaFold 3.0.1 was employed to generate structural predictions, evaluating monomeric or dimeric configurations (when the system has 2 genes). When possible, we annotated whether the novel systems were predicted as potentially defensive^69^ (GeneCLR hit in File S3). These integrated structural data were used to infer the presence of specialized antiviral effector or sensor domains by comparing the predicted folds to known defence-related protein architectures.

Genes of unknown functions were scored by both the ESM-650M_DF_ and the geneCLR_DF_ scores as defined in^69^. The methods were run on representative sequences when clustering the database at 95% of identity. Thus, we mapped the genes of unknown function to their representative 95 and retrieved the corresponding scores.

### Detection of homologs of P4/P2 defence genes in bacterial and bacteriophage genomes

We collected all defence-associated proteins in P4-like satellites and P2-like genomes, and then used this set with Diamond BlastP to search for homologs in the complete RefSeq bacterial genomes, including their plasmids. We used Diamond BlastP with the parameters --query-cover 50, --ultra-sensitive and --max-target-seqs 0. We then retrieved all matches with at least 85% identity with the query sequence and consider these as homologs of the matched defence genes. We did a similar analysis to search for homologs of P4 or P2 defence-associated proteins in the dataset of 27022 bacteriophage genomes described above.

The genomic context of the homologs detected in bacterial genomes is assigned as a plasmid (if in a secondary replicon of the bacterial genome), a prophage (if in a region detected as such by geNomad), a satellite (if in a region detected as such by SatelliteFinder), a phage-plasmid (if the plasmid replicon correspond to known phage-plasmids^37^ or if geNomad detects a secondary replicon as a prophage) or an integrative mobile element (i.e., Integrative Conjugative Elements – ICEs –, Integrative Mobilizable Elements – IMEs – or iOriTs, if in a region detected as described in^34^). If the defence homolog was detected in a bacterial genomic region that does not correspond to any of the categories above, we assigned it as an “uncharacteristic genomic region”.

To account for the possibility that homology of defence systems is caused by vertical descent within the element, i.e., cases where an homolog of a P4/P2 defence gene was found in a bacterial region that corresponds to that same P4/P2, we calculated the core (minus the integrase) wGRR (see section above) between all P4 and P4*, and all P2 and P2*, and discarded homologous matches with elements that had a wGRR of at least 0.95.

### Phylogenetic analyses

To compute the phylogenies of P4 and P2 elements, we extracted all their respective core proteins (except for the integrase), and aligned each set of proteins individually, using using mafft-linsi^70^ (v. 7.490, default parameters). The individual alignments of core proteins were then concatenated in two alignments (one with the P4 core proteins, another with the P2 core proteins). For elements where some core genes were missing, a sequence of gaps (“-“) was added. The resulting alignment was trimmed with clipkit^71^ (v. 1.3.0, default parameters). We then used IQ-Tree^72^ (v. 1.6.12) to build the phylogenetic trees, with the options –bb 1000 to run the ultrafast bootstrap option with 1000 replicates and -m TEST option for automatic model choice. The resulting tree files were visualized and edited using the v5 webserver of iTOL^73^

The phylogenies of individual defence genes were inferred by aggregating in single fasta files all the protein sequences corresponding to homologous matches of specific defence genes (PD-T7-2, Lit and Avs_V). Systems with more than one gene (i.e., PD-T7-2) were aggregated in separate files, and each gene was processed separately. The alignments and phylogenies were assembled and visualized as described above.

### Functional annotation of uncharacteristic genomic regions

Homologous matches of defence genes of P4/P2 in bacterial chromosomic regions that do not correspond to known MGEs were defined as “uncharacteristic genomic regions”. To analyse the genomic context of these homolog genes, we extracted 20 genes upstream and 20 genes downstream. These regions were then further annotated with DefenseFinder^19^ and PADLOC^20^, P4 and P2-specific HMM markers, and EggNogMapper^62^, all with the same parameters described above in each section. We also used PseudoFinder^25^, as described above, to detect pseudogenes in these uncharacteristic genomic regions, and the intergenic pseudogenes detected were further classified with P4 and P2 specific HMM profiles. To maintain specificity, we excluded from the search for core markers the integrase of P4/P2, as well as Delta and AlpA of P4, since homologs to these components can be found in other MGEs.

### Visualization of genomic regions

Genome visualization across the manuscript uses Clinker version 0.0.3^74^ with the -i parameter set at 0.3 at the other parameters were left as default. The JSON output files were edited to better represent the colors of the gene clusters.

## Acknowledgements

We thank all the Microbial Evolutionary Genomics Unit members for the regular discussions throughout the project. The authors also wish to acknowledge the students of the 2025 M2 Microbiology course at Institut Pasteur (Pierric Chuppa, Hugo Cibla, Léa Dunand-Sauthier, Esteban Francois, Thibault Frisch, Theana Gardais, Chiara Gouviaux-Deletraz, Dorian Joffres, Ekaterina Korotaeva, Kevin Koule, Jeane Le Floch, Nicolas Le Jalle, Laly Meyer, Laura Ochsenbein, Orane Pion, Maël Rochefort, Corentin Rohel, Alexandre Roubaud, Inès Santos and Sofia Tagkalidou), who collaborated in the preliminary assays of infection the candidate defence systems. This work used the computational and storage services (MAESTRO cluster) provided by the IT department at Institut Pasteur, Paris. We acknowledge support from the Laboratoire d’Excellence IBEID Integrative Biology of Emerging Infectious Diseases [ANR-10-LABX-62-IBEID], the Agence Nationale de Recherche [ANR 23 CE20 0046 01 TRIADE] and [ANR-20-CE12-0004 TransfoConflict] and the HORIZON programme [HORIZON-MSCA-2024-PF-01 TriSH]. This work used the computational and storage services (MAESTRO cluster) provided by the IT department at Institut Pasteur, Paris.

