## Supplementary Figures for "Shuttling, swapping and mixing: the rapid modular evolution of antiviral repertoires in temperate phages and their satellites"

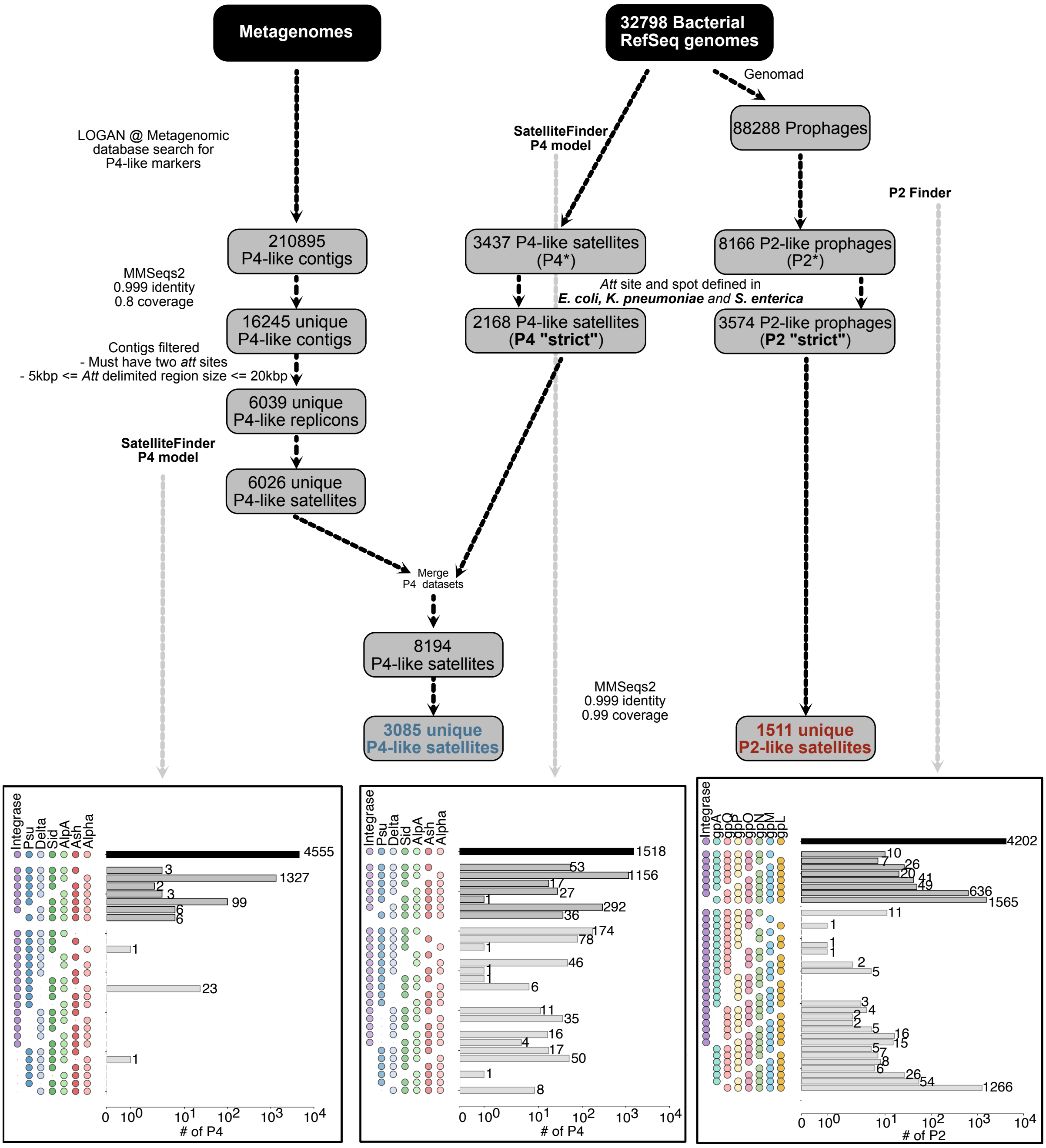


**Figure S1. The dataset of P4-like satellites and P2-like prophages used in this study.** Schematic visualization of the original datasets (in black ellipses) and the processing (filtering, de-replication) that led to the final datasets of 3085 P4-like satellites and 1511 P2-like prophages. Shown are also the distribution and classification (in Types A, B, or C, corresponding to their completeness) of the total number of P4s and P2s detected before filtering by att sites in the 3 main species, as well as the total number of unique P4-like satellites detected from metagenomic data, before merging with the P4 detected from RefSeq genomes. In these barplots, the x-axis shows the number of elements (in log scale), and the y-axis shows the possible combinations of core genes detected (Type A, all core genes were found; Type B, one of the core genes is missing or undetected; Type C, two of the core genes are missing or undetected).

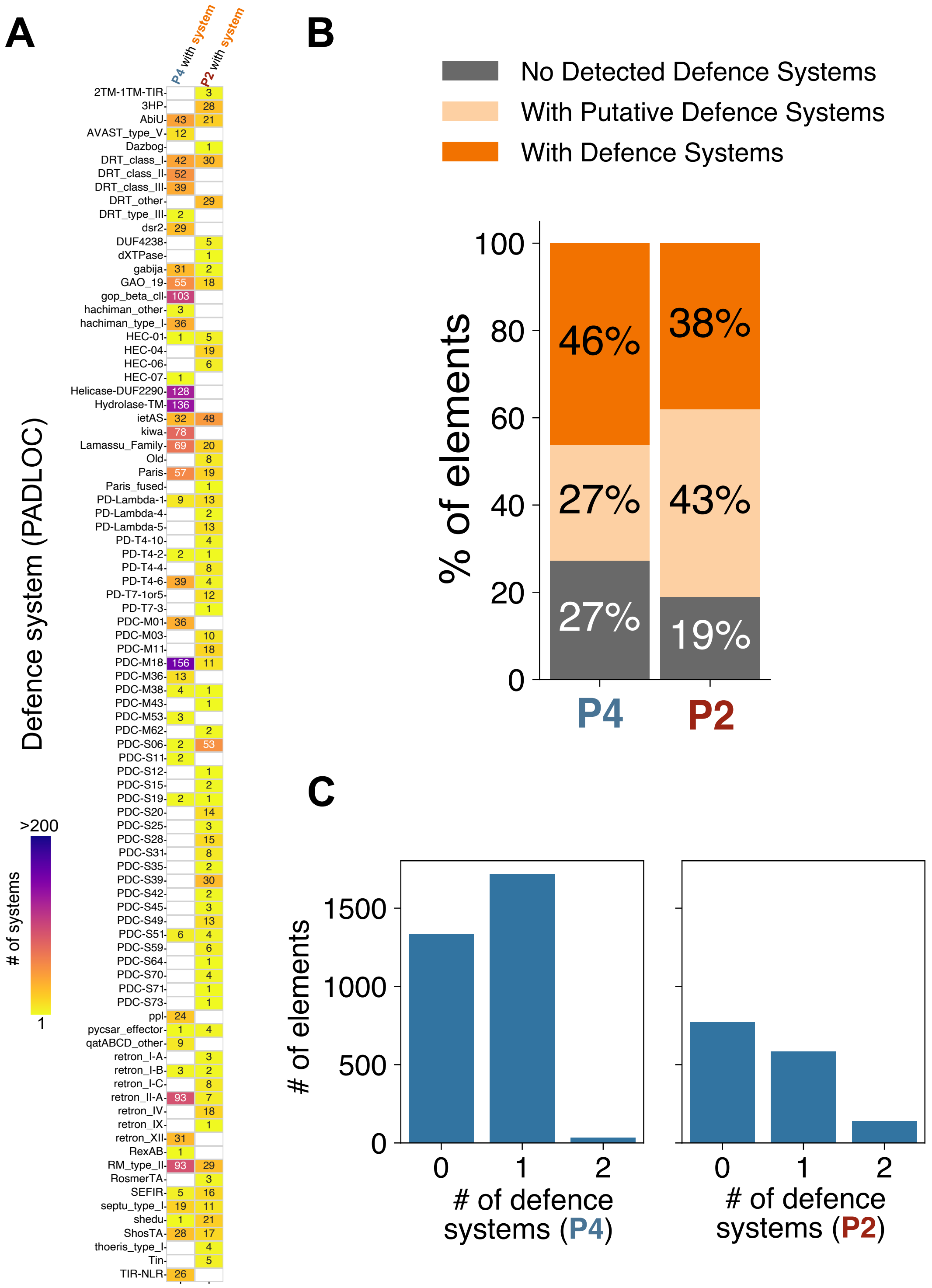


**Figure S2. Contribution of PADLOC to the diversity of defence systems detected in P4s and P2s and distribution of defence systems per element. A.** In the y-axis are the different types of complete (two leftmost columns) and putative (two rightmost columns) defence systems detected by PADLOC in P4 (1st column) and P2 (2nd column) elements. The number of elements with a given defence system is shown in each position of the heatmap, with darker colours corresponding to systems more frequently detected. **B.** The proportion of P4 and P2 elements with complete defence systems (orange) as detected by DefenseFinder only, as well as the putative defence systems (light orange) detected by DefenseFinder. **C.** The distribution of the number of complete defence systems per element, for P4 and P2, respectively.

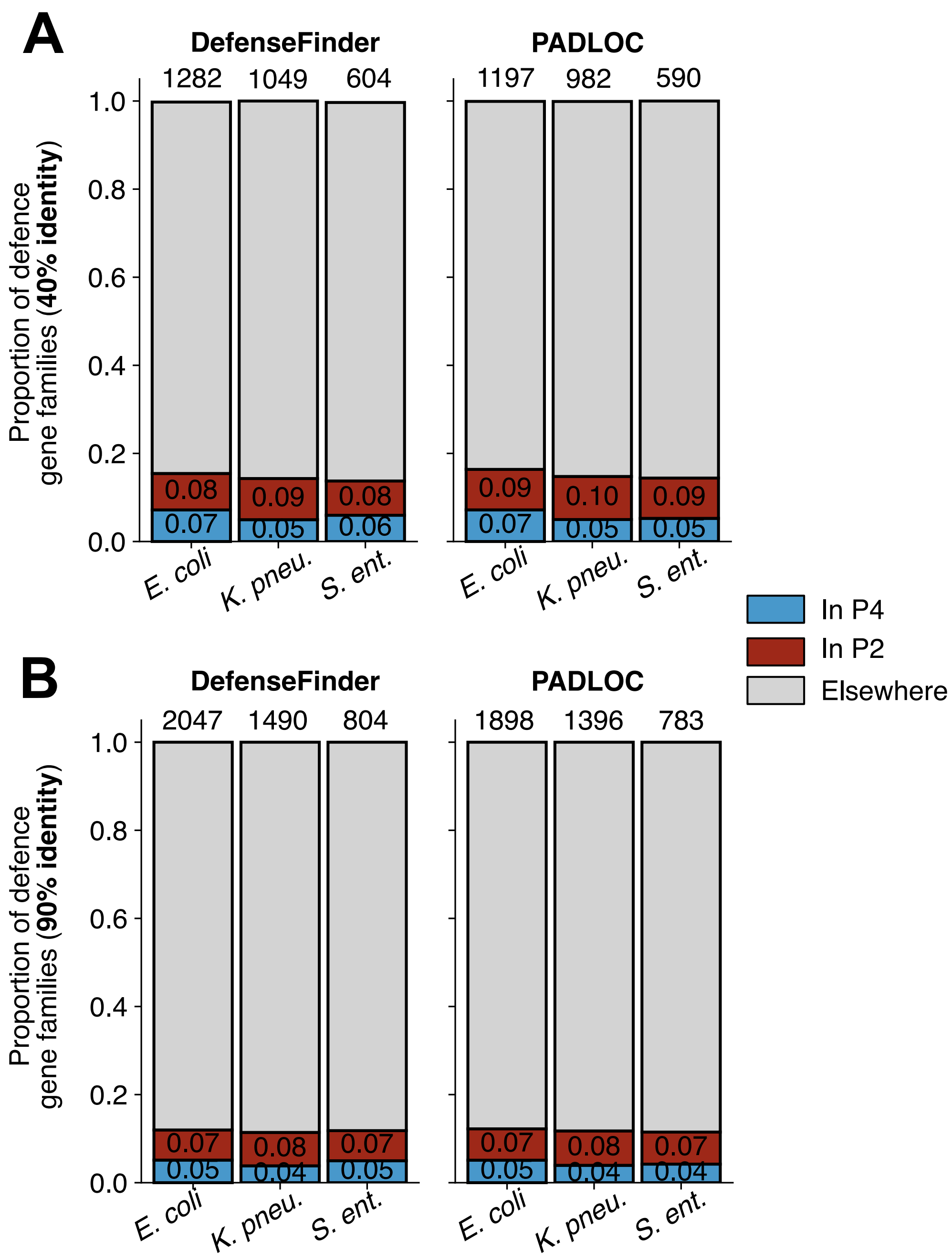


**Figure S3. Contribution of P4 and P2 to the pan-defences of their bacterial hosts.** Defence genes were inferred using DefenseFinder and PADLOC in the bacterial genomes that encode P4 and/or P2 for each of the three focal species, independently. **A**. Genes from complete systems detected by Defense Finder (left) or PADLOC (right), were clustered at 40% identity. From these gene families, the stacked barplots represent the proportion of gene families that include at least one gene from one P4 (in blue), at least one gene from one P2 (in red), or if the gene family was never found within one of these two elements (grey). At the top of each bar is shown the total number of defence gene families for each species, in accordance with each defence detection software. A residual number of gene families is found both in P4 and P2 (bars not visible) **B.** Same as in **A**, but with gene families assembled by clustering the defence genes at 90% identity. At this identity level, no defence gene families are found in both P4 and P2.

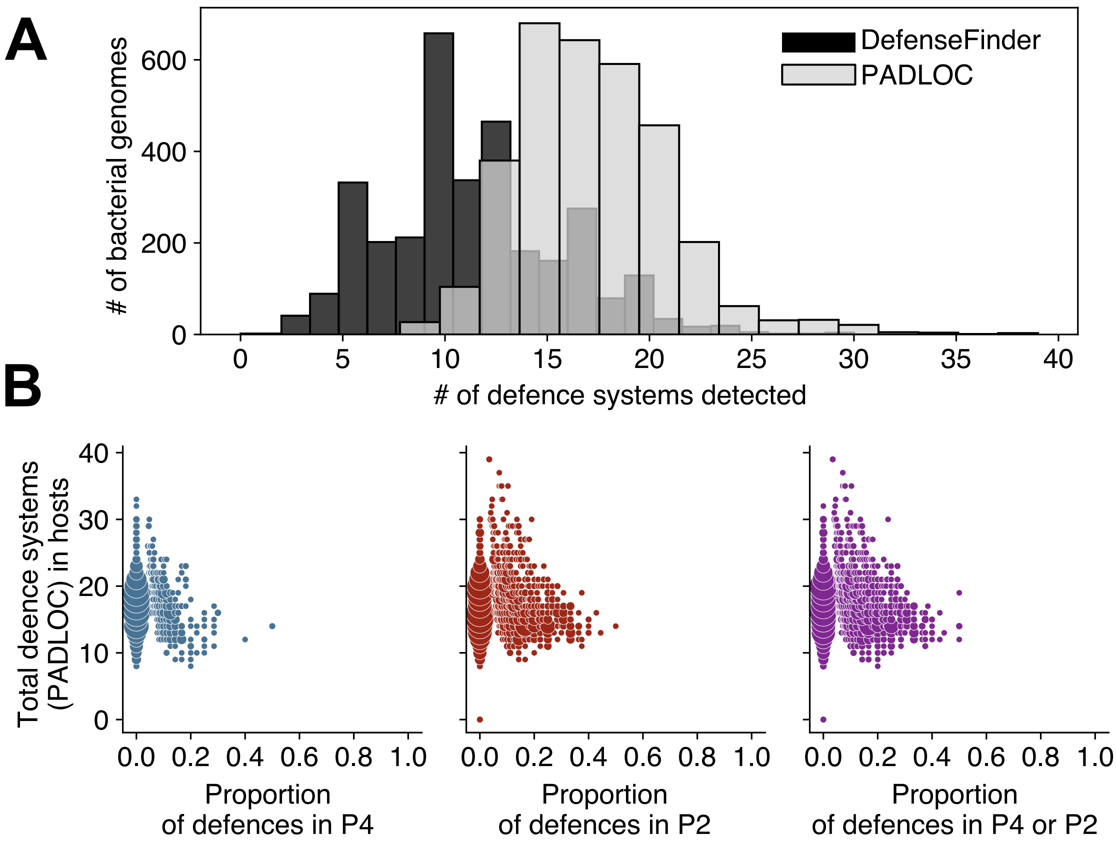


**Figure S4. Defence systems detected in bacterial genomes. A**. Distributions of the number of defence systems per bacterial genome, according to DefenseFinder (black bars) and PADLOC (grey bars). **B.** The proportions of PADLOC defence systems encoded by P4 in hosts that have at least one P4 (leftmost panel), encoded by P2 in hosts that have at least one P2 (middle panel), or encoded either by P4 or P2 in hosts that have at least one of either element (rightmost panel) are shown in the x-axis, with the total number of defence systems detected in their hosts displayed in the y-axis. The size of the circles correspond to the number of cases for each (x,y) pair.

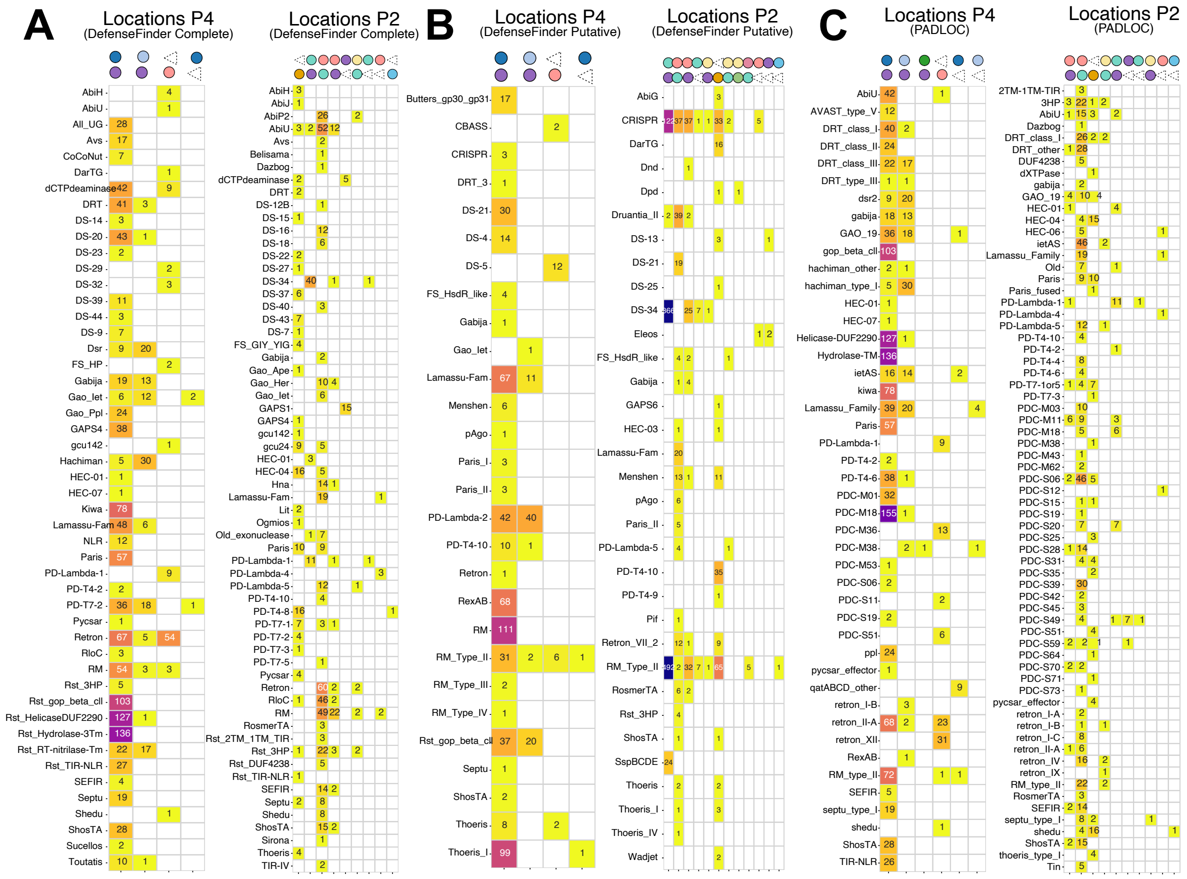


**Figure S5. Distribution of defence systems across hotspots in P4 and P2 genomes. A.** Distribution of complete systems detected by DefenseFinder. **B.** Distribution of putative defense systems detected by DefenseFinder. **C.** Distribution of defence systems detected by PADLOC. In all three panels, the number of elements where a given defence system is found at a specific location is shown in each position of the heatmap, with darker colours corresponding to systems more frequently detected. The combinations of two markers at the top of each column correspond to the core genes (or the att sites, indicated as triangles) that flank the defence system (e.g., in A, the first column corresponds to the spot between Integrase and Psu).

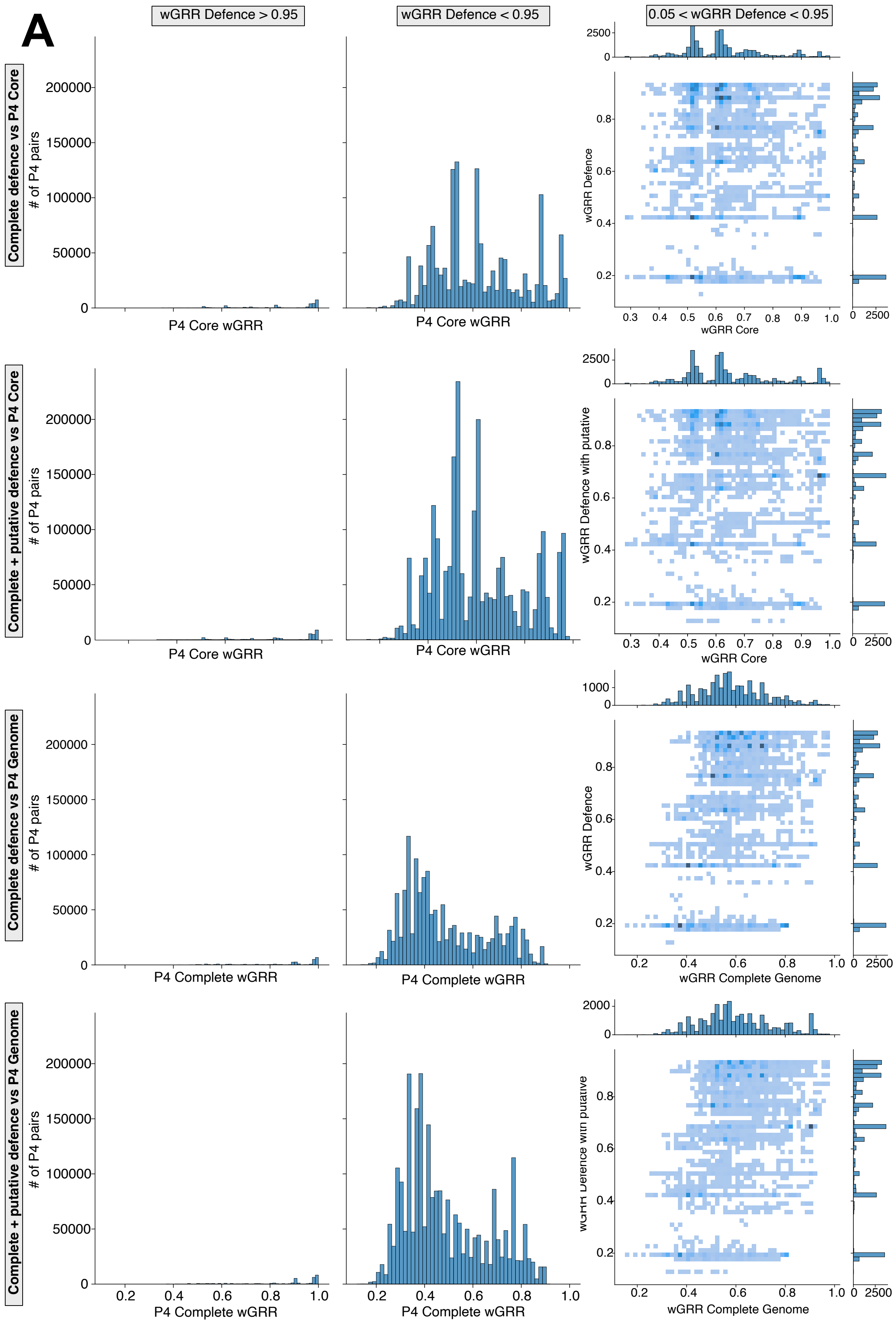
A
wGRR Defence > 0.95
wGRR Defence < 0.95
0.05 < wGRR Defence < 0.95
Complete defence vs P4 Core
Complete + putative defence vs P4 Core
Complete defence vs P4 Genome
Complete + putative defence vs P4 Genome
### of P4 pairs
P4 Core wGRR
P4 Complete wGRR
wGRR Core
wGRR Complete Genome
wGRR Defence
wGRR Defence with putative

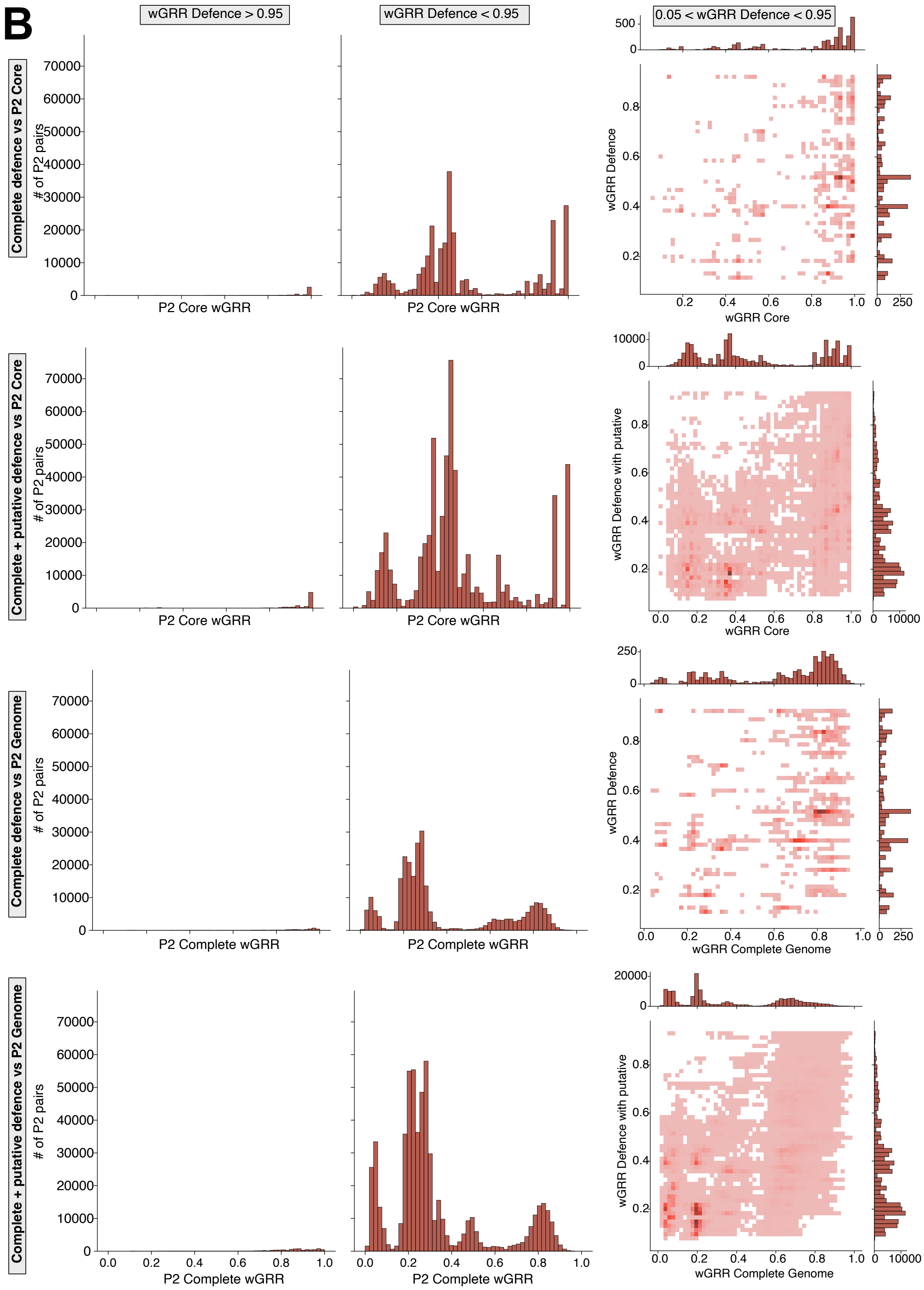
B
wGRR Defence > 0.95
wGRR Defence < 0.95
0.05 < wGRR Defence < 0.95
Complete defence vs P2 Core
Complete + putative defence vs P2 Core
Complete defence vs P2 Genome
Complete + putative defence vs P2 Genome
### of P2 pairs
P2 Core wGRR
P2 Complete wGRR
wGRR Defence
wGRR Defence with putative
wGRR Core
wGRR Complete Genome

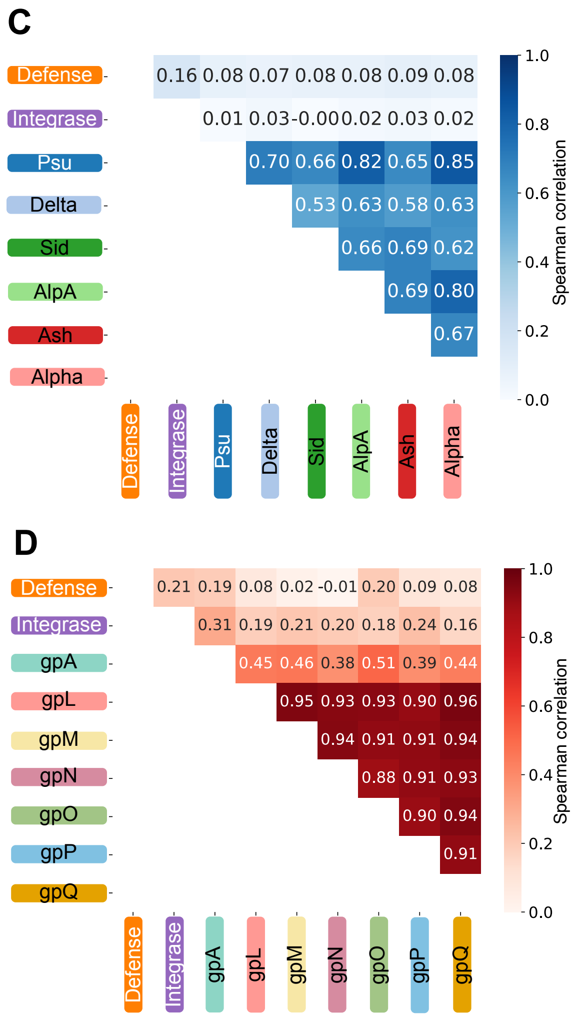


**Figure S6. Relationship between antiviral repertoires and the core genome in pairs of P4-like satellites and in pairs of P2-like prophages. A.** Left, histogram with the distribution of counts for pairs of P4 genomes with similar defence genes (wGRR defence at least 0.95, first and third row taking into account only complete defence systems, second and fourth rows taking into account both complete and putative defence systems), across all the range of genomic similarity of the elements (x-axis, core genes in first and second row, full genome in third and fourth row). Center, histogram with the distribution of counts for pairs of P4 genomes with distinct defence genes (wGRR defence at most 0.05), across all the range of genomic similarity (x-axis). Right, a matrix representing bin-based joint distribution of the defence similarity (y-axis) and genomic similarity (x-axis) of pairs of P4, for the defence similarity values that are not comprised by the left and center distributions (i.e, 0.05<wGRR defence <0.95). Stronger shades of color represent more populated bins. The marginal distributions in each axis represent the distribution of the univariate distribution of defence similarity (wGRR defence, y-axis) and P4 genome similarity (wGRR core/full, x-axis). **B.** Same as in **A**, but for P2 genomes. **C.** Spearman correlation values between the wGRR of defence genes, including both complete and putative systems, and the BlastP values of individual core genes of P4-like satellites, for the set of all pairs of P4s. Colors of a stronger shade indicate higher Spearman correlation values. **D.** Same as in **C**, but for the defence and core genes of P2.

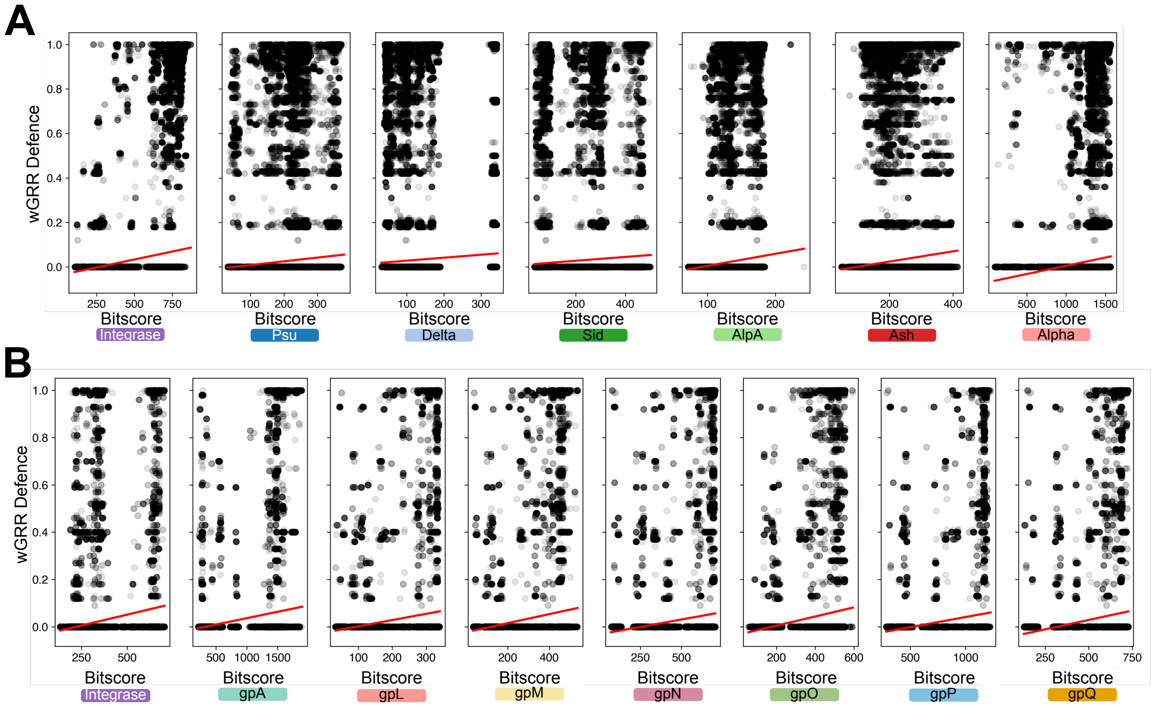


**Figure S7. Detailed correlation between defence genes and individual core genes in pairs of P4 and pairs of P2. A.** Each panel corresponds to the correlation between the wGRR of defence genes and the similarity (BlastP) of specific core genes of P4, across all pairs of P4s. The color of the dots corresponds to the frequency of combinations of values (darker colors represent more frequent cases). **B.** Same as in **A**, but for pairs of P2s.

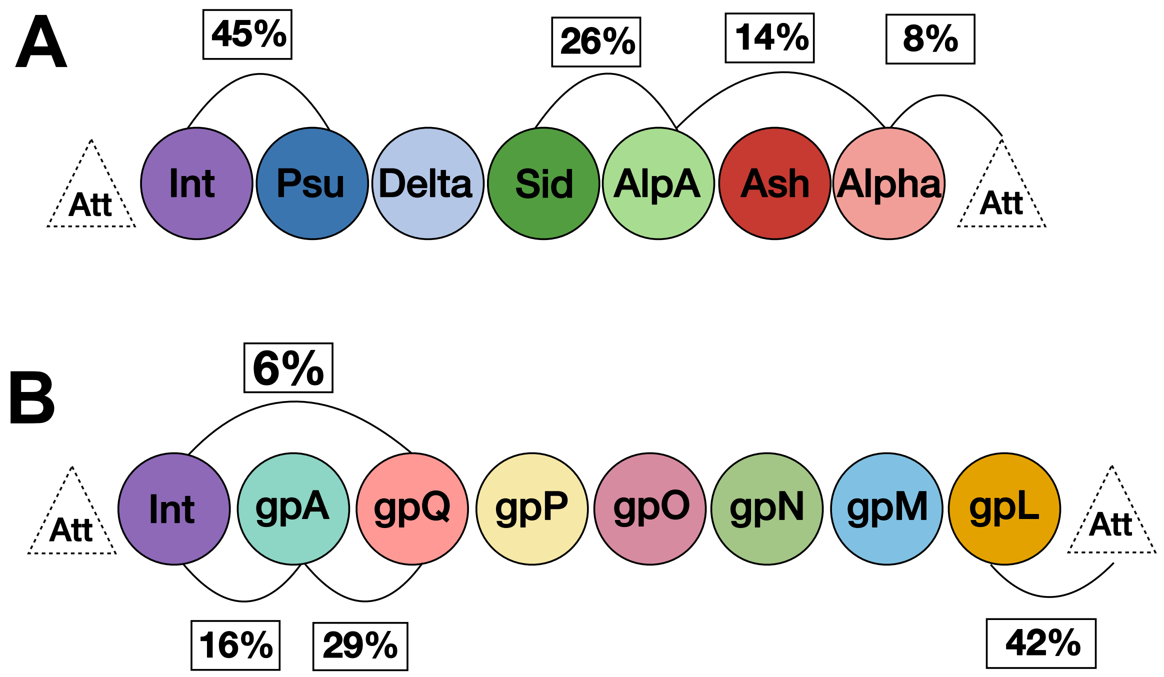


**Figure S8. Hotspots of intergenic pseudogenes.** Edges between circles (core genes) indicate the proportion of intergenic pseudogenes detected at those locations. **A.** Main locations of intergenic pseudogenes of P4. **B.** Main locations of intergenic pseudogenes of P2.

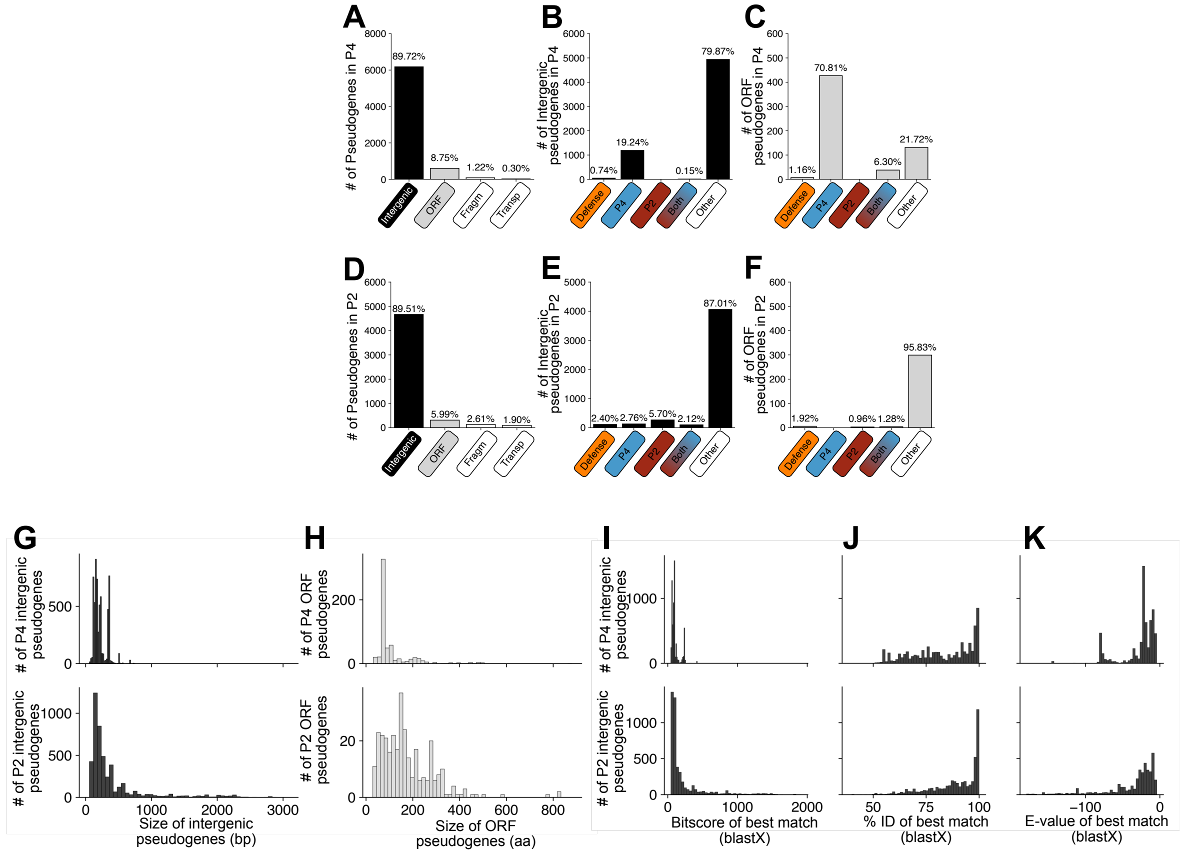


**Figure S9. Detailed characterization of pseudogenes detected in P4 and P2. A and D**. The quantification of the type of pseudogene detected in P4 and P2. **B and E**. Quantification of intergenic segments proposed as pseudogenes, whose most similar complete proteins are homologous to antiviral-associated profiles, P4 markers, P2 markers or both P4 and P2 markers (i.e., integrase). **C and F**. Quantification of ORF proposed as pseudogenes that are homologous to antiviral-associated profiles, P4 markers, P2 markers or both P4 and P2 markers (i.e., integrase). **G.** Nucleotide size distribution of the intergenic sequences identified as pseudogenes in P4 (**top**) or P2 (**bottom**). **H.** Aminoacid size distribution of the ORFs that are predicted as pseudogenes in P4 (**top**) or P2 (**bottom**). **I, J and K.** Distribution of the bitscore, %Id and e-value for the RefSeq genes that are best matches for the intergenic pseudogenes in P4 (**top**) or P2 (**bottom**).

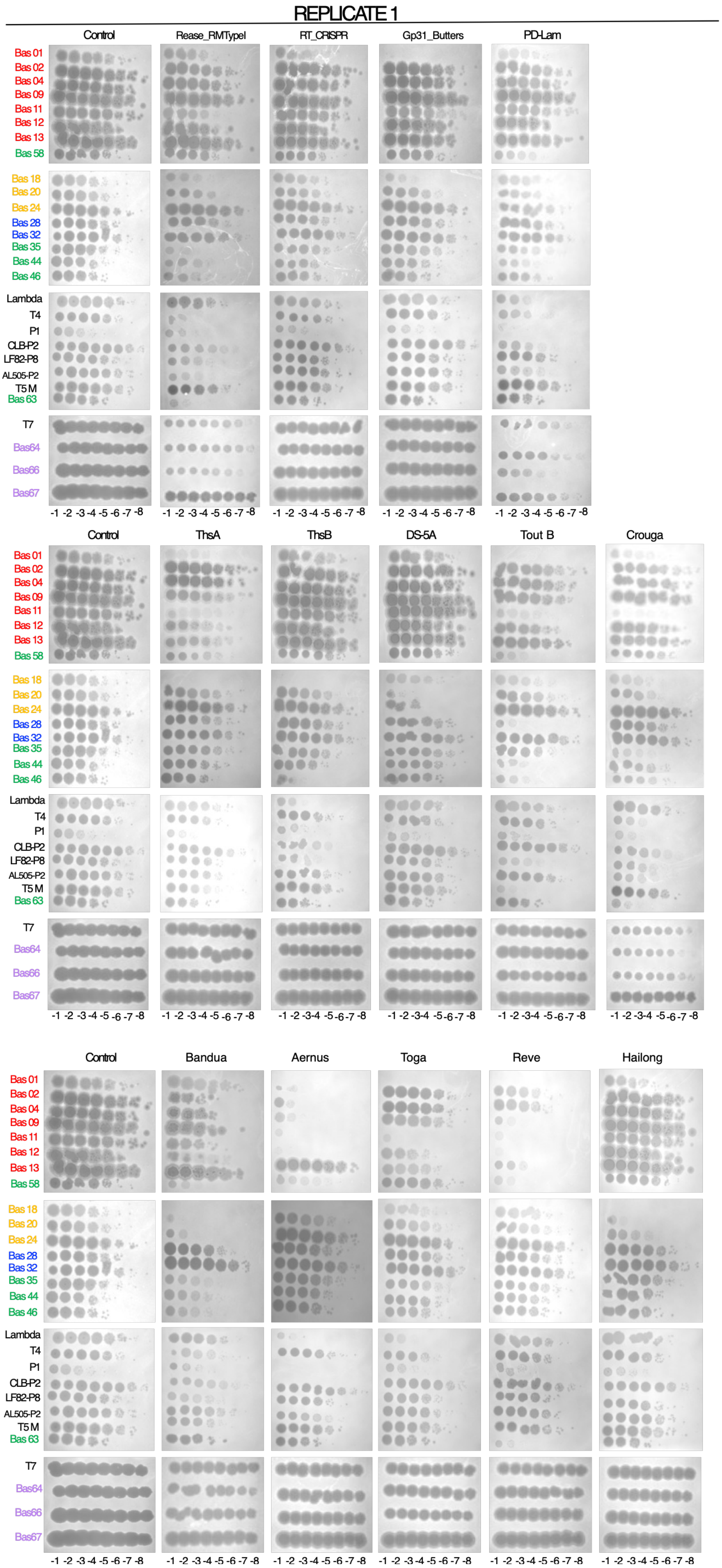
REPLICATE 1
Control
Rease_RMTypeI
RT_CRISPR
Gp31_Butters
PD-Lam
ThsA
ThsB
DS-5A
Tout B
Crouga
Bandua
Aernus
Toga
Reve
Hailong
Bas 01
Bas 02
Bas 04
Bas 09
Bas 11
Bas 12
Bas 13
Bas 58
Bas 18
Bas 20
Bas 24
Bas 28
Bas 32
Bas 35
Bas 44
Bas 46
Lambda
T4
P1
CLB-P2
LF82-P8
AL505-P2
T5 M
Bas 63
T7
Bas64
Bas66
Bas67
-1 -2 -3 -4 -5 -6 -7 -8

### REPLICATE 2

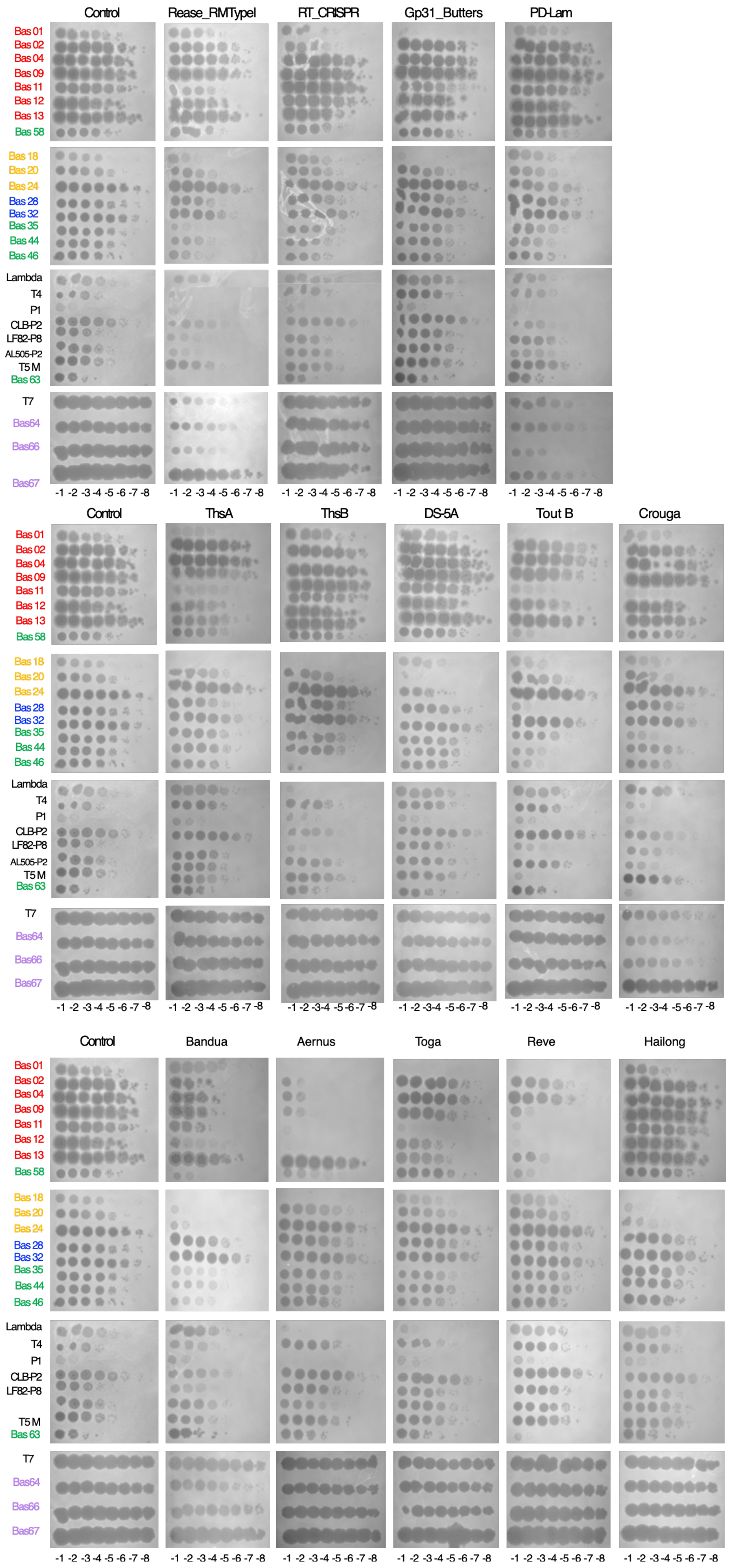
Control
Rease_RMTypeI
RT_CRISPR
Gp31_Butters
PD-Lam
Bas 01
Bas 02
Bas 04
Bas 09
Bas 11
Bas 12
Bas 13
Bas 58
Bas 18
Bas 20
Bas 24
Bas 28
Bas 32
Bas 35
Bas 44
Bas 46
Lambda
T4
P1
CLB-P2
LF82-P8
AL505-P2
T5 M
Bas 63
T7
Bas64
Bas66
Bas67
-1 -2 -3 -4 -5 -6 -7 -8
Control
ThsA
ThsB
DS-5A
Tout B
Crouga
Control
Bandua
Aernus
Toga
Reve
Hailong

**Figure S10**. **Anti-phage activities of the 11 putative and 8 unknown defence systems against a panel of 28 phages**. Columns correspond to the different putative and novel defence systems, whilst rows correspond to the different phages used in the infections. The first column in either replicate represent infection of strains with the control (empty) plasmid. The two independent replicates for the infection assays are shown. Tenfold serial dilutions of high titer phage lysates were spotted on E. coli strains carrying either the control plasmid (pFR66) or a plasmid expressing the defense system, in the presence of aTc to induce expression. Plaque-forming units were measured for each phage from these representative images corresponding to two biological replicates.

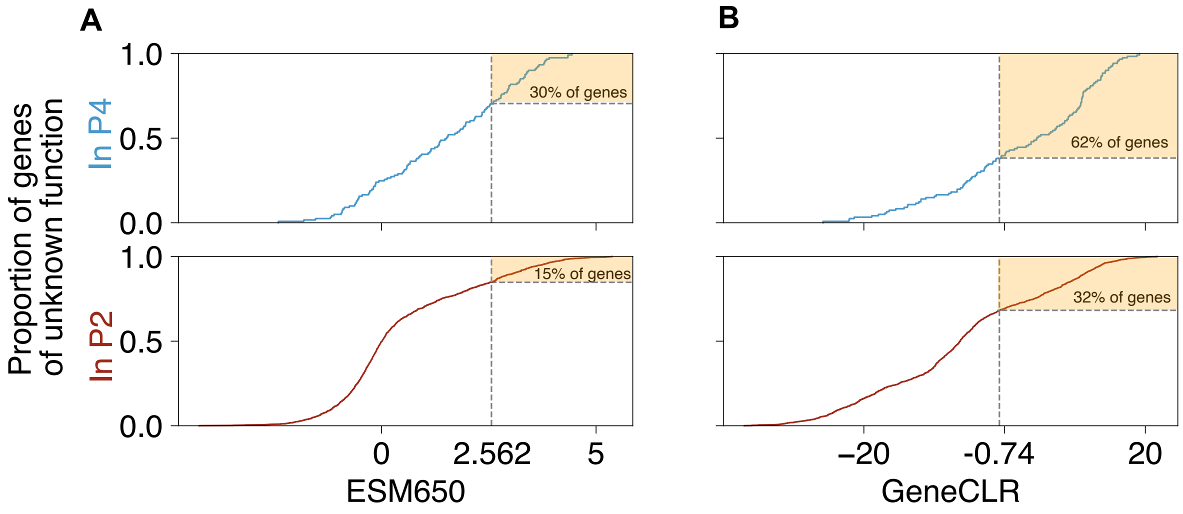


**Figure S11**. **Cumulative distribution of the "defence scores" in genes of unknown function**. Genes of unknown function in P4 (limited to RefSeq genomes, top) and P2 (bottom), were analysed using two different computational methods: ESM650, for the analysis of protein domains (left); and GeneCLR, which takes both protein domains and gene context into account (right). The critical values of significance for each method (above which proteins are predicted to have a defence function) are indicated as grey dashed vertical lines in each plot (2.562 for ESM650, and -0.74 for GeneCLR, as defined in 10.1101/2025.01.08.631966).

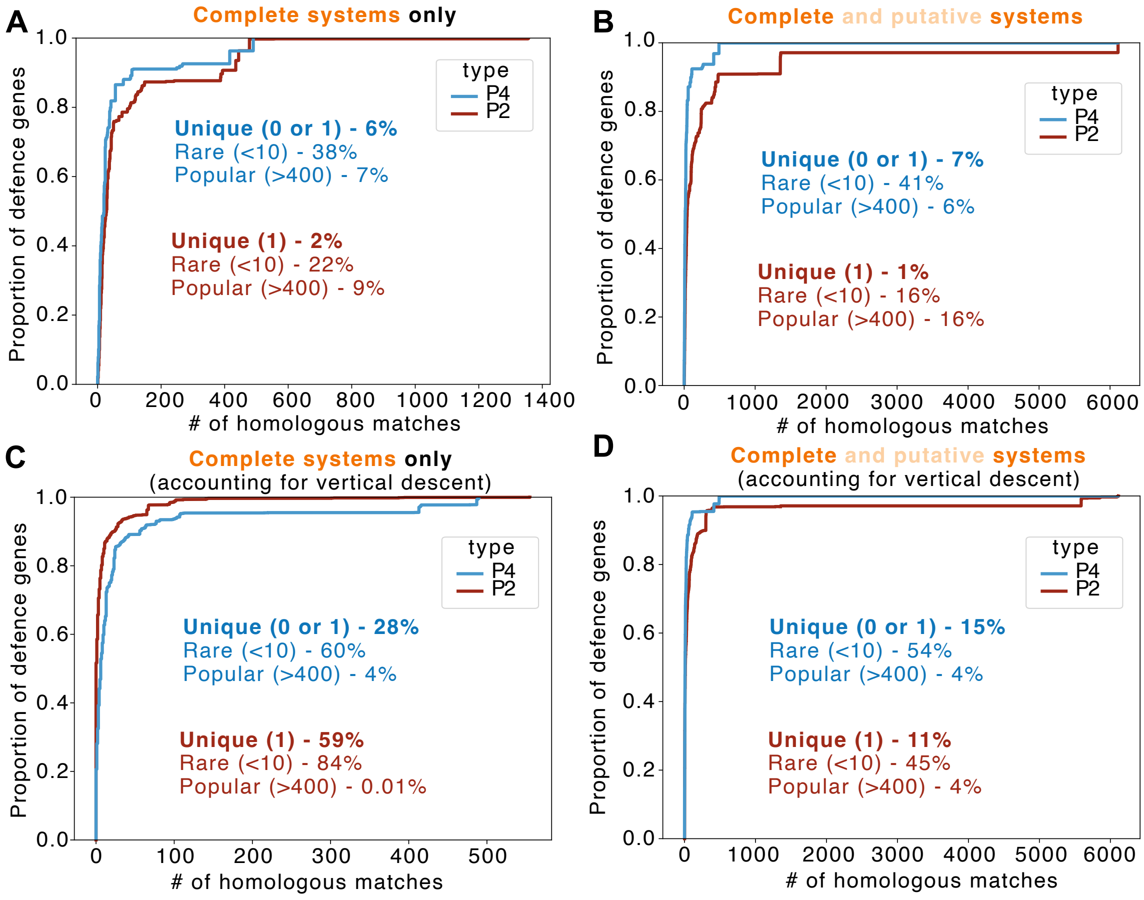


**Figure S12. Empirical cumulative distribution functions of homologous defence matches of P4 and P2.** In the y-axis, the proportion of defence genes with that has the number of homologous matches in complete bacterial genomes, that are shown in the x-axis. In **A and B**, values are relative to the raw data from the Diamond BlastP output that fulfills the homology criteria (>85% identity). In **C and D**, the values are relative to the matches after a control for vertical descent (i.e., if the homologous protein was detected in a region of the bacterial genomes that contains a P4 or P2 with a wGRR>=0.95 relative to the P4 or P2 that encoded the queried defence gene). In **A and C**, only defence genes from complete defence systems are queried for homologs. In **B and D**, defence genes from both complete and putative systems are queried for homologs.

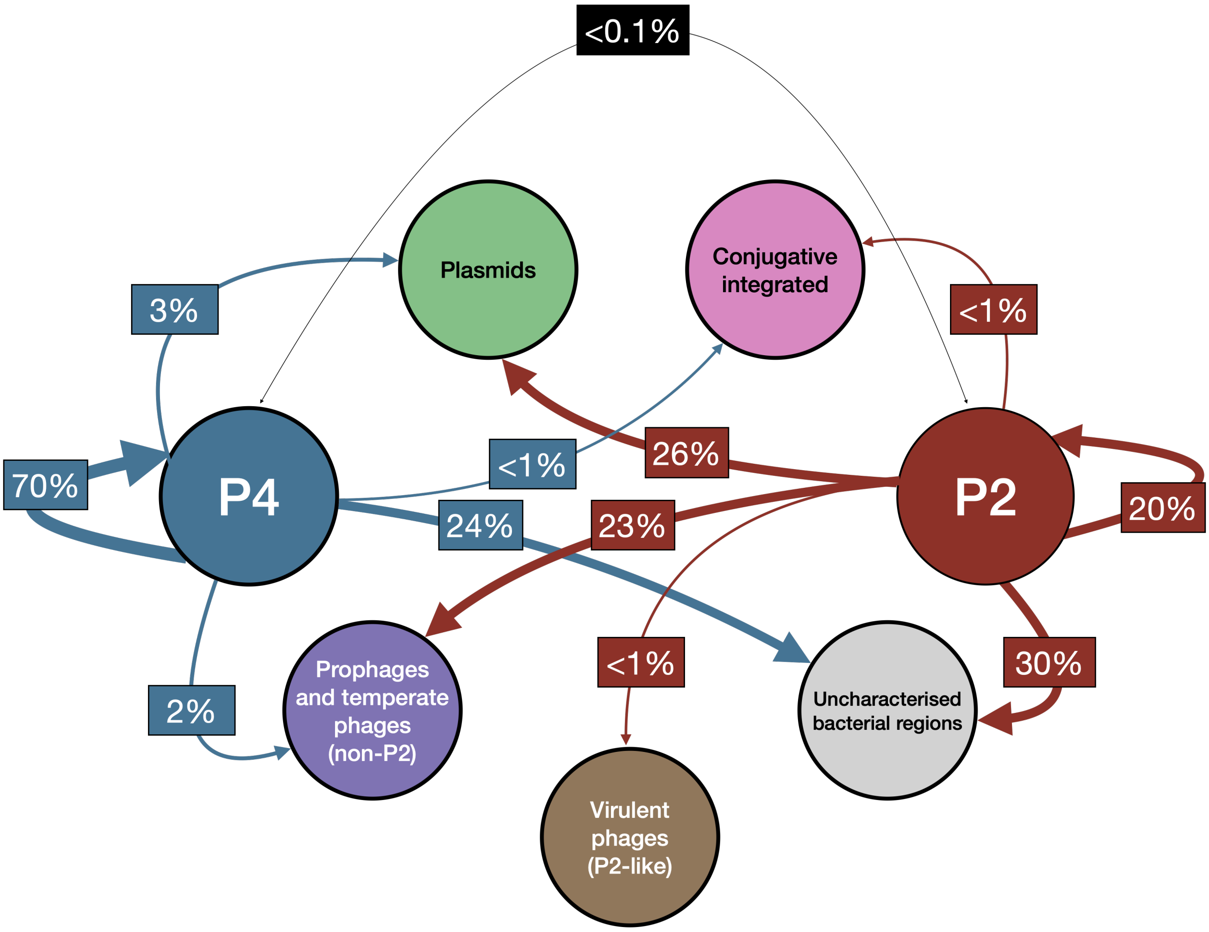


**Figure S13. Network of homology of defence genes from complete and putative systems with other bacterial genomic regions.**

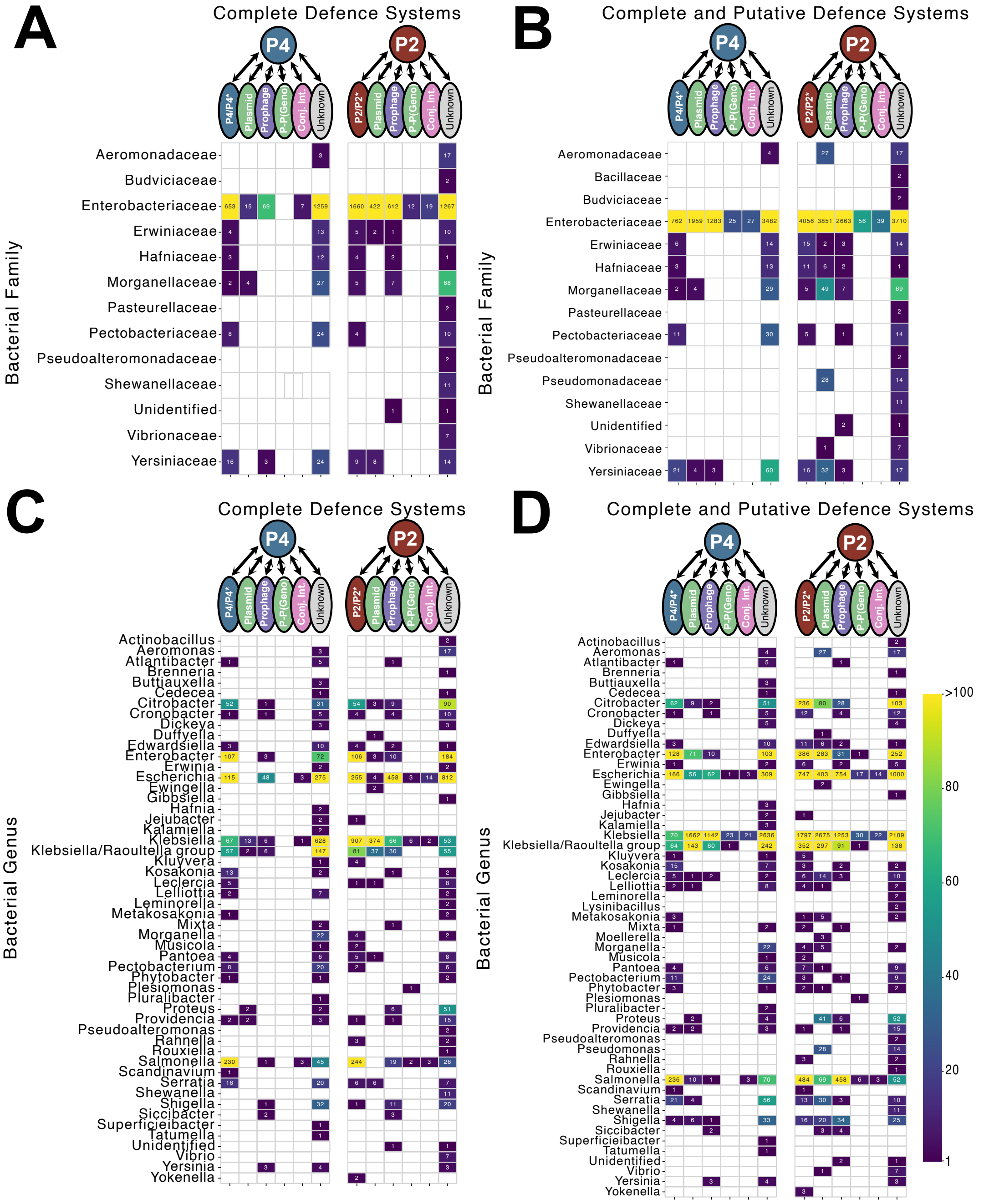


**Figure S14. Detection of homologs of defence genes of P4 and P2 across bacterial host families and genera. A.** The different bacterial families (y-axis) with their corresponding number of unique genes detected as homologs of defence genes from complete defence systems in P4 (left matrix) and P2 (right matrix) are detected, in each genomic context (x-axis). Yellow colors represent more unique genes detected as homologs, blue colors represent a lower number of unique genes detected as homologs, and white cells represent the lack of any homologs found for the host/genomic context combination. **B.** Same as in **A**, but considering defence genes from both complete and putative systems. **C and D**. Same as in **A** and **B**, but considering bacterial genera instead of families.

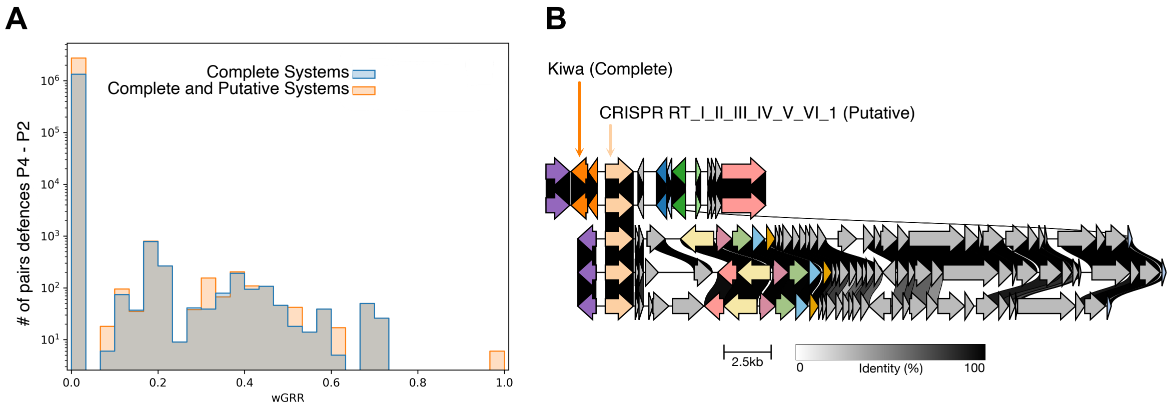


**Figure S15. Defence homologs are very rare between P4 and P2 genomes. A.** wGRR (x-axis) distribution for P4 and P2 pairs of defence systems (complete, blue bars, and complete and putative, orange bars). **B.** Unique example detected of a putative defense system shared between P4 and P2 genomes with high homology.

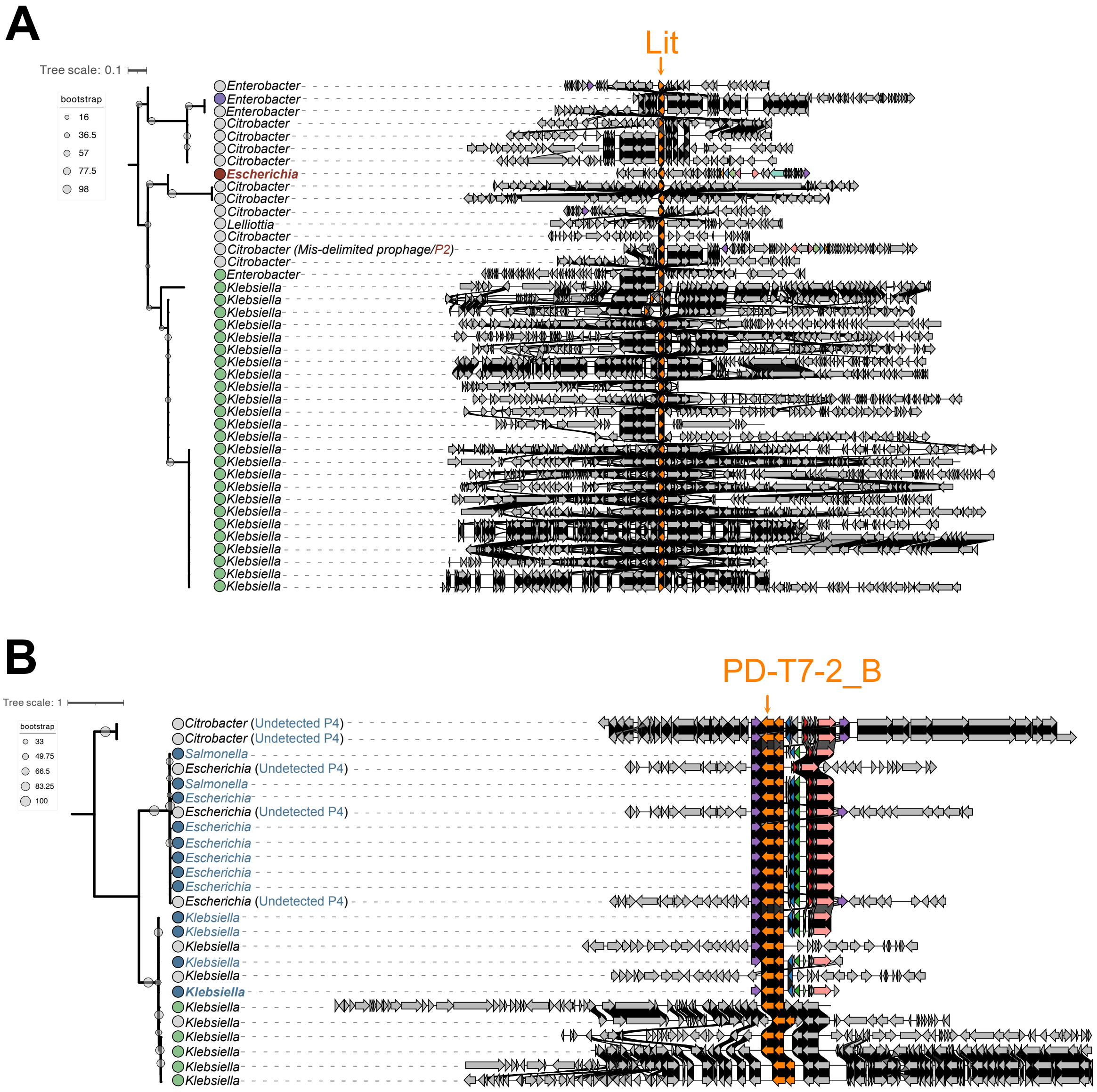


**Figure S16. Additional phylogenies of defence genes in P4 and P2 with homologs in dissimilar genomic contexts. A.** Phylogeny of the Lit gene, comprised of the gene found as a defence system in a focal P2 in Escherichia, and its homologs detected in either other P2s (red circles at the tree leaves), plasmids (green circles), prophages (violet circles) or uncharacterized genomic regions (grey circles). The genes corresponding to each of the elements (or to the segments that include the uncharacterized genomic regions) are shown in front of the tree, with larger plasmids truncated for visibility. **B.** Phylogeny of the PD-T7-2-B gene, comprised of the gene found in a PD-T7-2 defence system in a focal P4 in Klebsiella, and its homologs detected in either other P4s (blue circles at the tree leaves), plasmids (green circles) or uncharacterized genomic regions (grey circles). The genes corresponding to each of the elements (or to the segments that include the uncharacterized genomic regions) are shown in front of the tree, with larger plasmids truncated for visibility.

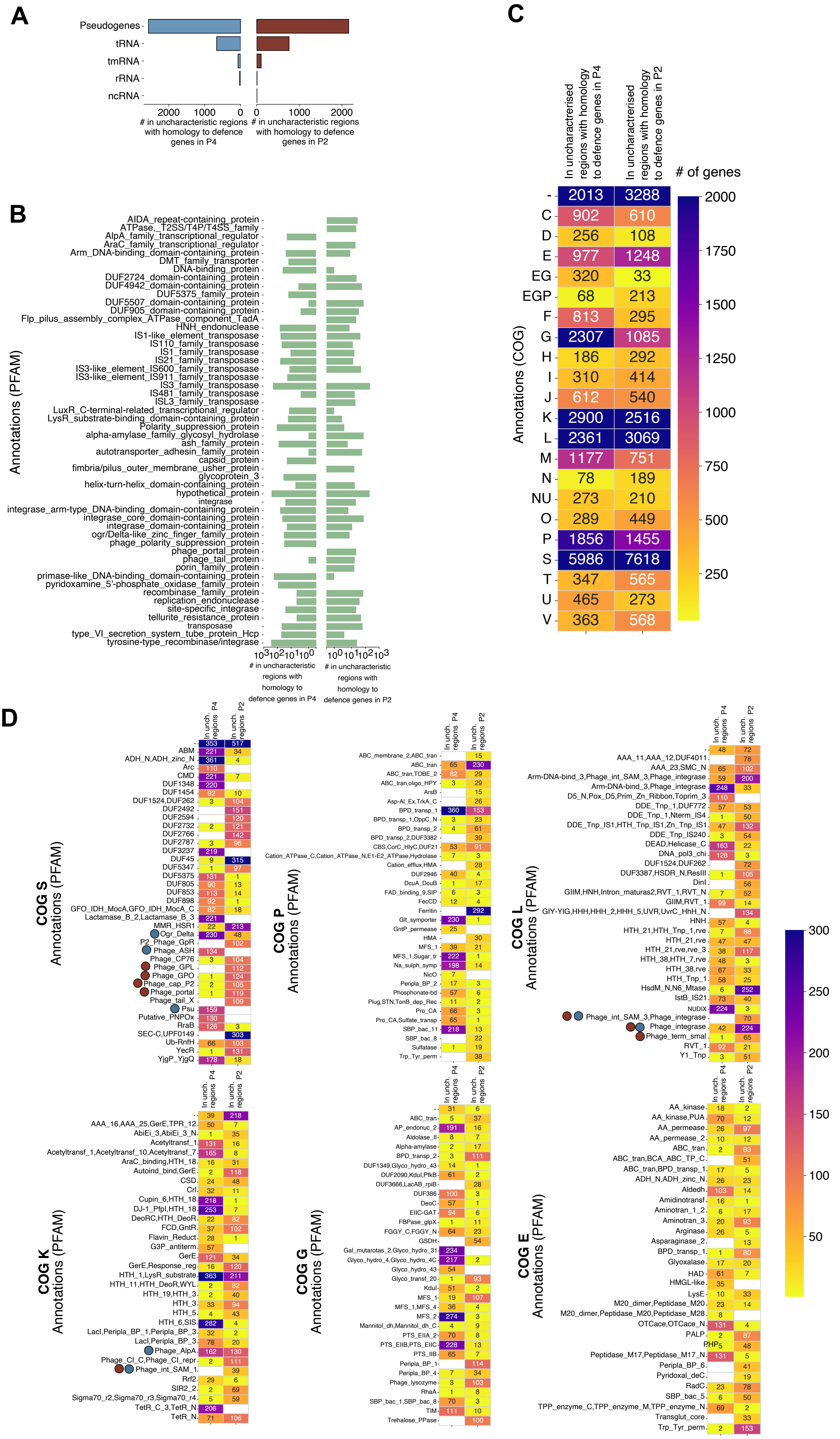
A
Pseudogenes
tRNA
tmRNA
rRNA
ncRNA
### in uncharacteristic regions with homology to defence genes in P4
### in uncharacteristic regions with homology to defence genes in P2
B
Annotations (PFAM)
### in uncharacteristic regions with homology to defence genes in P4
### in uncharacteristic regions with homology to defence genes in P2
C
In uncharacterised regions with homology to defence genes in P4
In uncharacterised regions with homology to defence genes in P2
### of genes
Annotations (COG)
- 2013 3288
C 902 610
D 256 108
E 977 1248
EG 320 33
EGP 68 213
F 813 295
G 2307 1085
H 186 292
I 310 414
J 612 540
K 2900 2516
L 2361 3069
M 1177 751
N 78 189
NU 273 210
O 289 449
P 1856 1455
S 5986 7618
T 347 565
U 465 273
V 363 568
D
COG S Annotations (PFAM)
COG P Annotations (PFAM)
COG L Annotations (PFAM)
COG K Annotations (PFAM)
COG G Annotations (PFAM)
COG E Annotations (PFAM)
In unch. regions P4
In unch. regions P2

**Figure S17. Functional annotation of uncharacterized genomic regions with homologs to defence genes in P4 and P2. A.** At the center, the circular bar plot showing the relative proportions of the different main COG categories that are associated with unknown genomic regions with homologs to defence genes of P4 (blue bars) and P2 (red bars). The most abundant categories are expanded in the surrounding barplots, which show the relative proportions of the PFAM terms associated with unknown genomic regions with homologs to defence genes of P4 (blue bars) and P2 (red bars). PFAMs that identify potential P4 or P2 cores genes are highlighted with blue or red arrows, respectively. **B.** Quantification of the genomic features other than ORFs that are detected in unknown genomic regions with homologs to defence genes in P4 (left bars) or P2 (right bars). These features were inferred from the NCBI annotations of bacterial genomes.

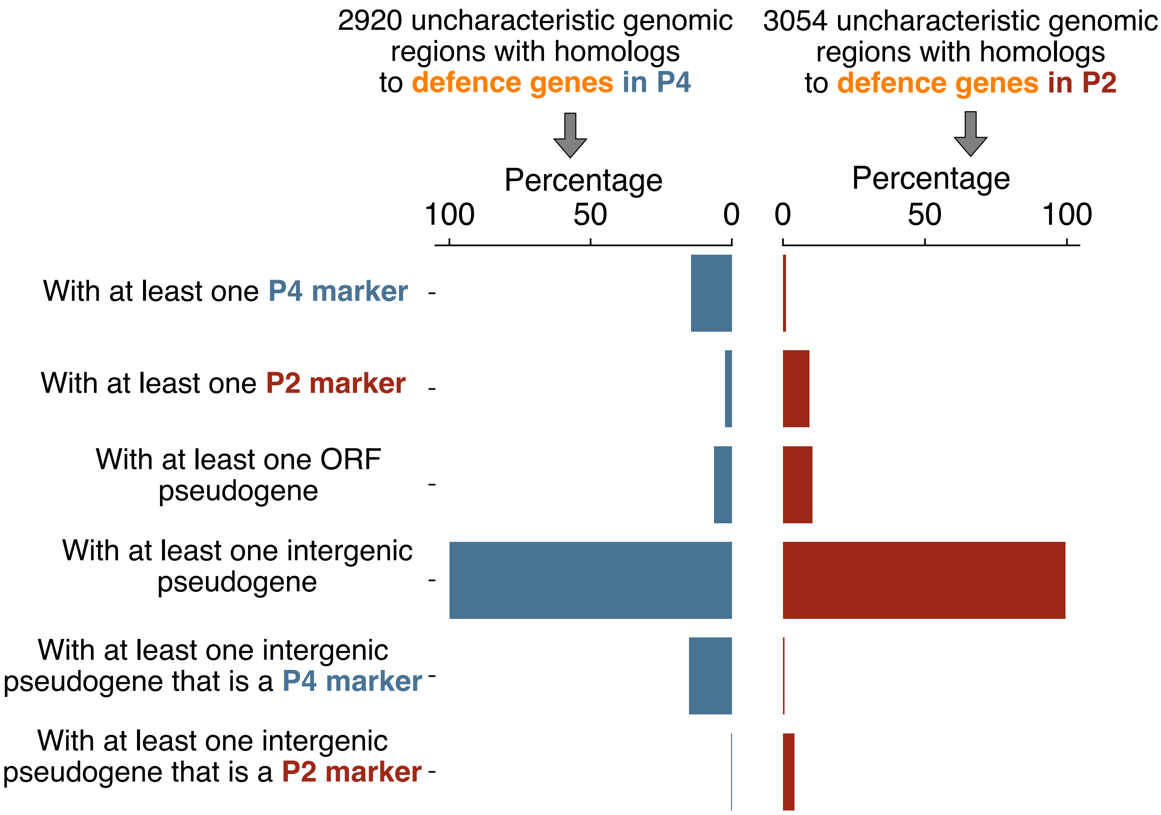


**Figure S18. Quantification of pseudogenes and P4-like/P2-like markers in unknown genomic regions with homologs to antiviral genes encoded by P4 and P2, including regions with homolgs to defence genes from putative defence systems in P4 and P2.** Characterization of uncharacterized genomic regions that encode homologs to defence genes, from both complete and putative systems, detected in P4 (left bars, blue) and uncharacterized genomic regions that encode homologs to defence genes, from both complete and putative systems, detected in P2 (right bars, red). We quantified the proportion of these regions that encode P4/P2 markers either as predicted ORFs (first two bars) or intergenic pseudogenes detected by PseudoFinder (last two bars).
