## Supplementary material for "Shuttling, swapping and mixing: the rapid modular evolution of antiviral repertoires in temperate phages and their satellites": FileS2_PutativeSystemsAnnotations.pdf

### REase::RM Type 1 (pFD408)

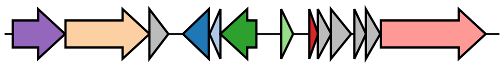

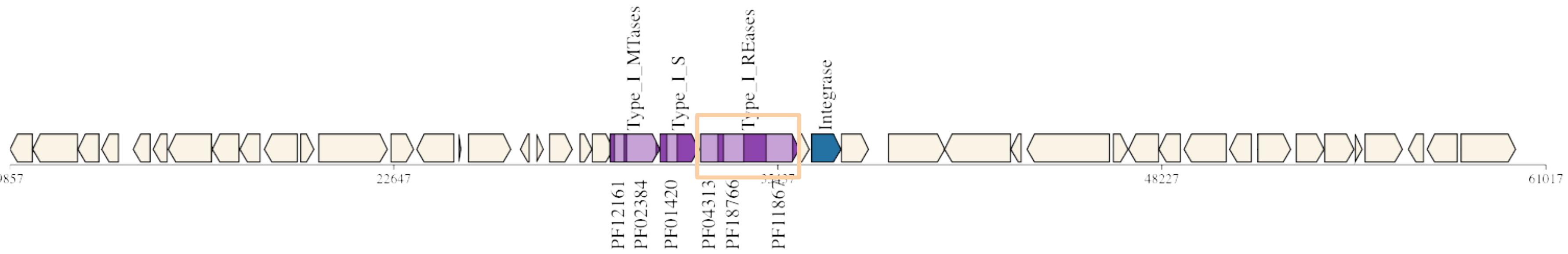


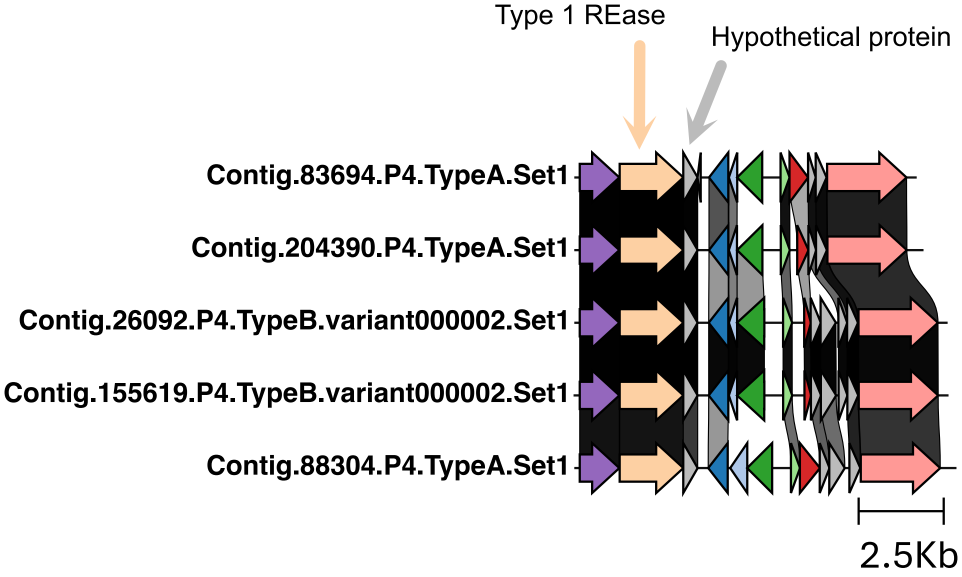


This putative system contains a REase of a RM Type 1, but is missing the other genes. It has an hypothetical protein next to it, that is conserved across P4 (along the REase gene)

#### RT::CRISPR (pFD410)

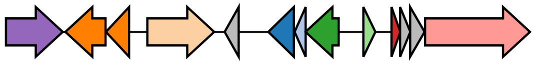

This partial system contains a reverse transcriptase associated with a CRISPR system. The domain could be part of a gene that has other functions. Of note, it is the only case of a defense gene (from partial or full systems) that is found recently shared between P4 and P2 (i.e., with high %identity). Unclear whether it requires other genes, as in P4 there is a small protein (in an opposite direction) and in P2 the neighbouring genes vary

**KLPN001.0523.00014.001C.P4.TypeB.variant0002.Set1**

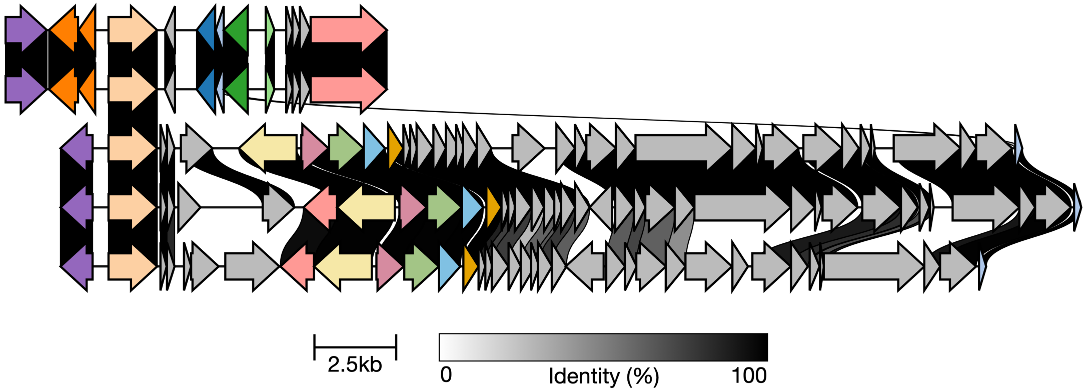

### GP31::Butters (pFD411)

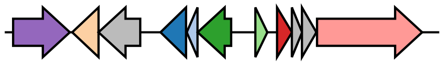

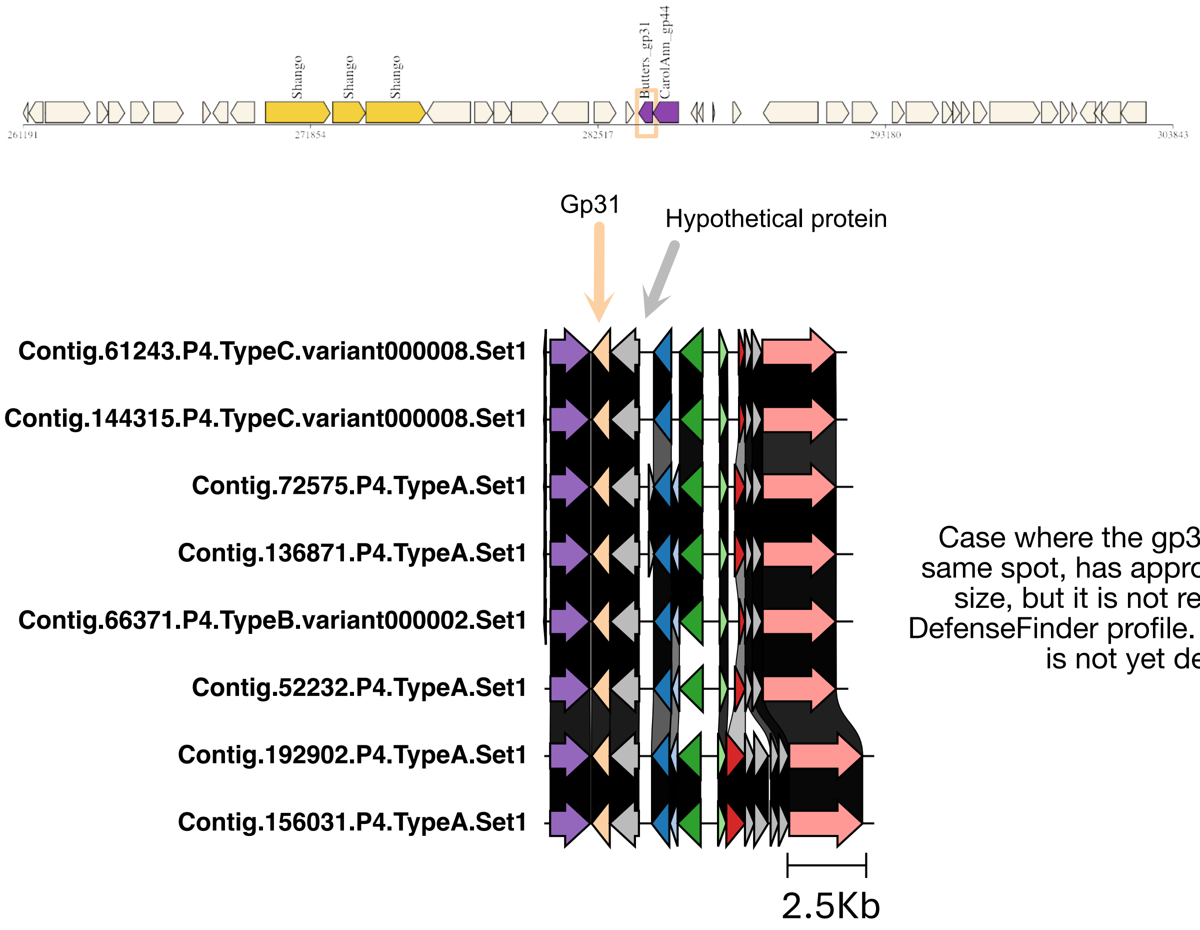


Case where the gp30 protein is in the same spot, has approximately the same size, but it is not recognised by the DefenseFinder profile. Likely a variant that is not yet described?

### PD_Lam_2C & PD_Lam_2B & cII ::PD Lambda 2 & Rst_gop_beta_cii (pFD413)

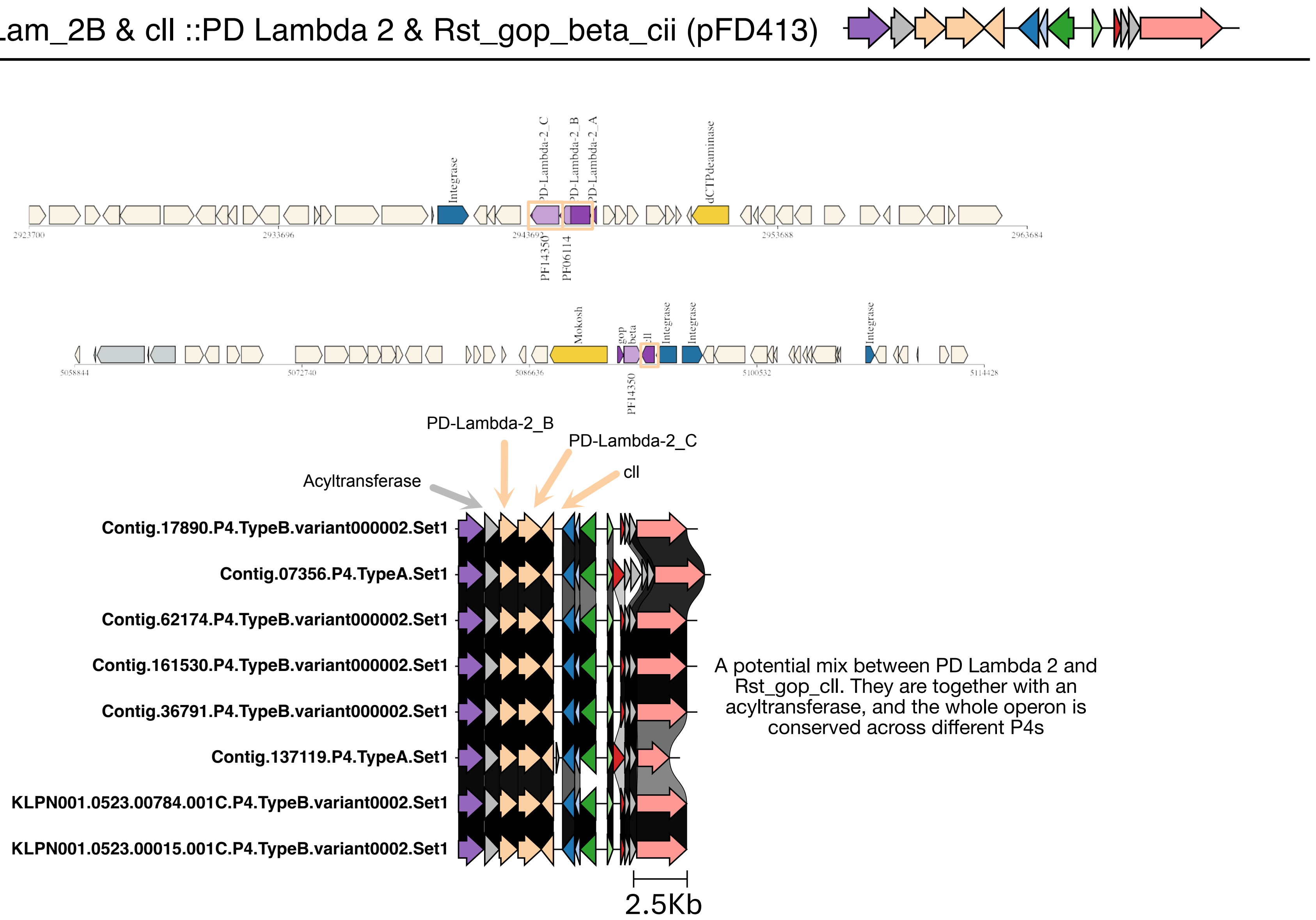


A potential mix between PD Lambda 2 and Rst_gop_cII. They are together with an acyltransferase, and the whole operon is conserved across different P4s

### ThsA::Thoeris (pfD414)

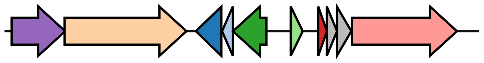

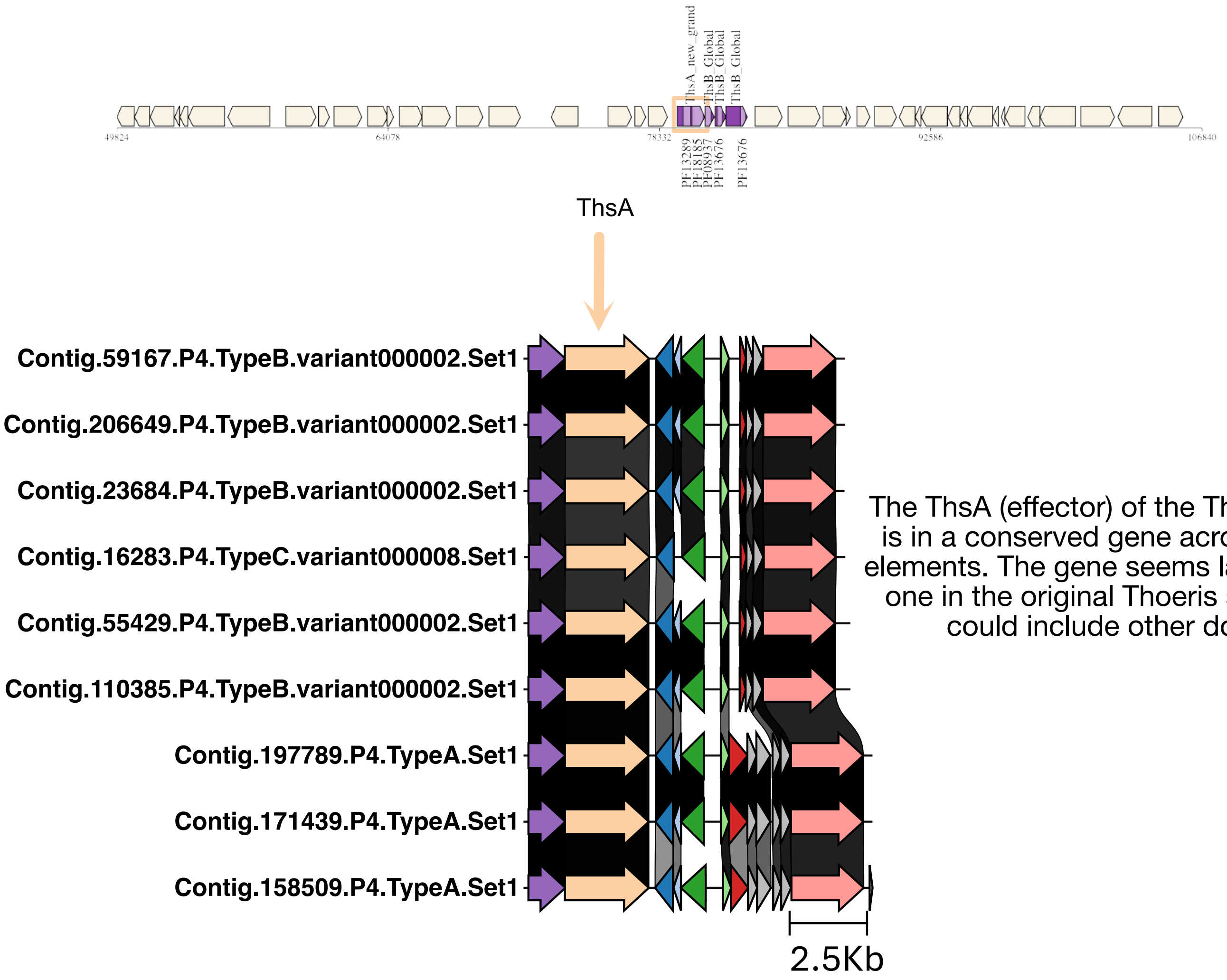


The ThsA (effector) of the Thoeris system is in a conserved gene across many P4 elements. The gene seems larger than the one in the original Thoeris system, and could include other domains.

### ThsB::Thoeris (pfD415)

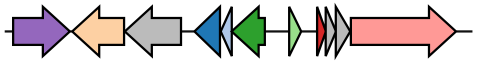

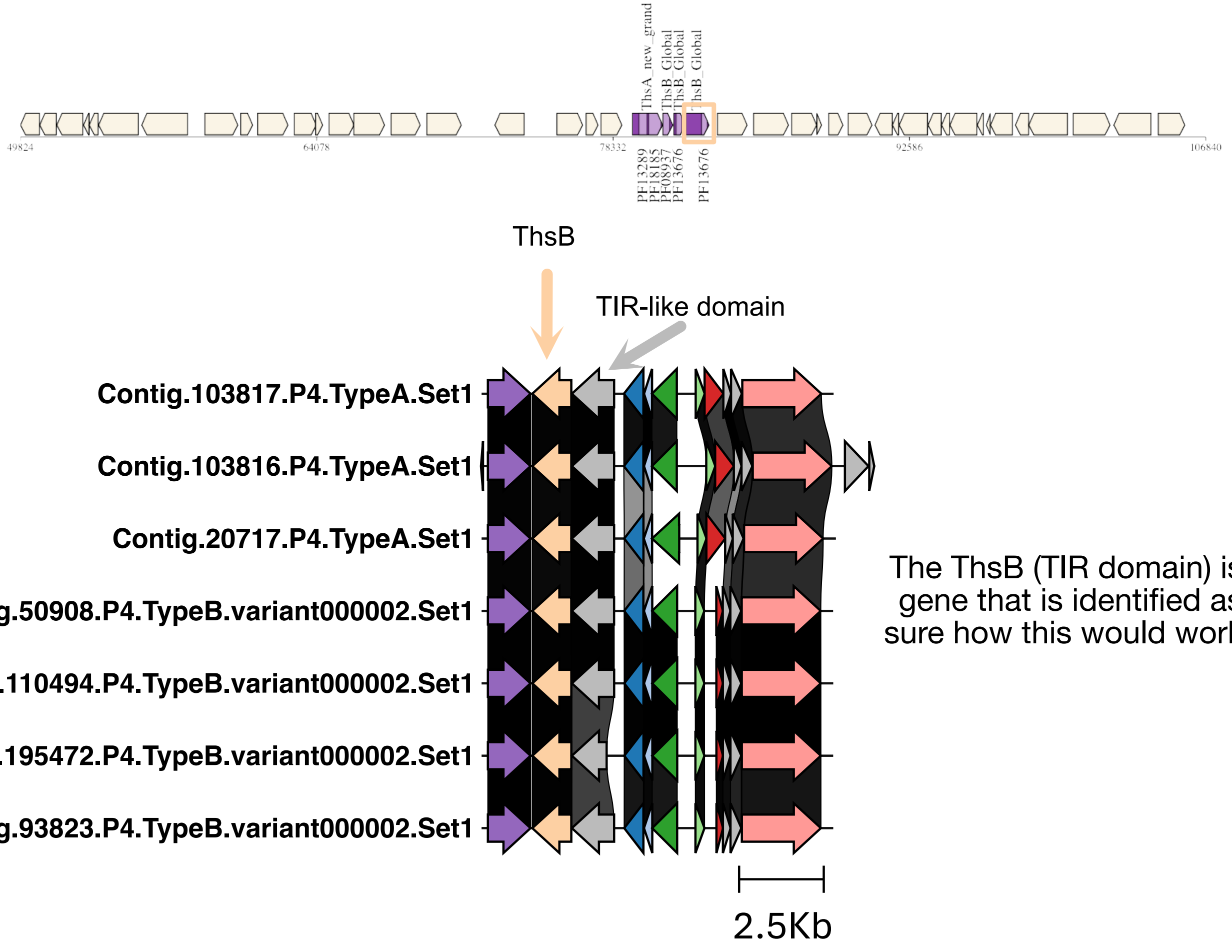


The ThsB (TIR domain) is found with another gene that is identified as a TIR domain. Not sure how this would work without an effector.

DS-5 A :: DS-5 (pfD416)

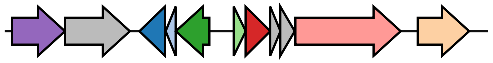

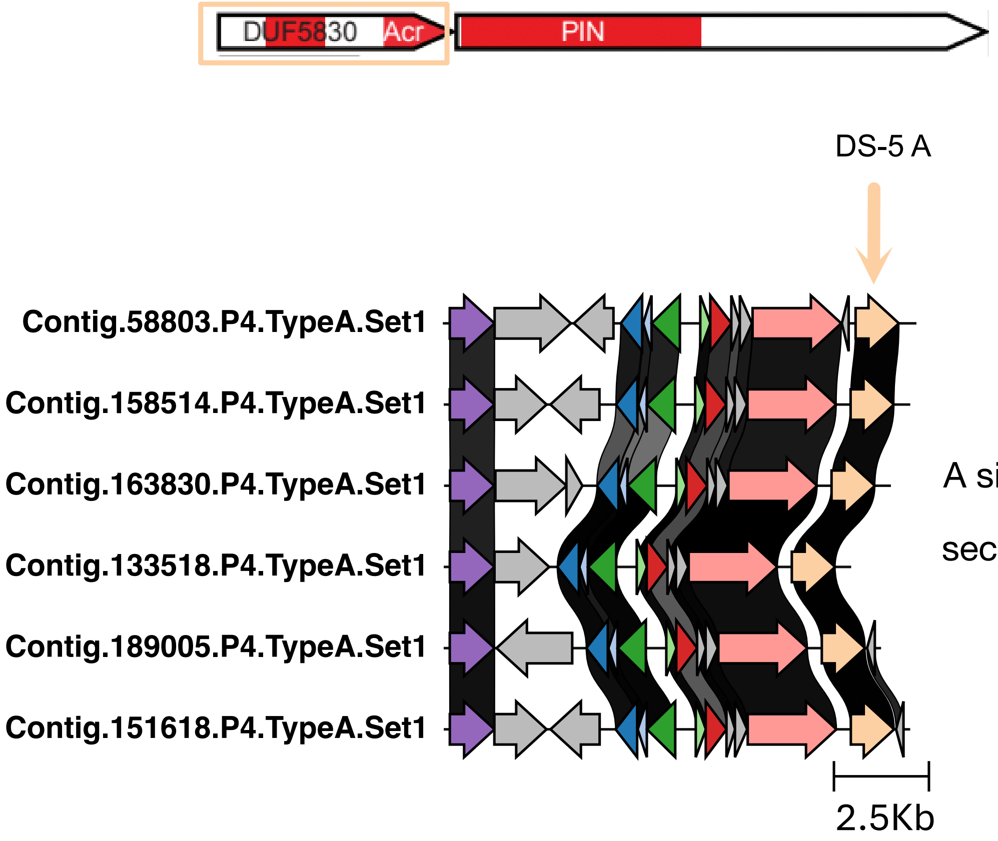


A single gene of the DS-5 system is found conserved in the secondary antiviral hotspot of P4.

### ToutB & PD-T4-10_A :: Toutatis & PD-T4-10_A (pfD417)

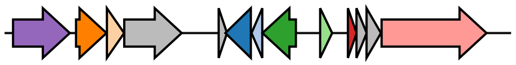

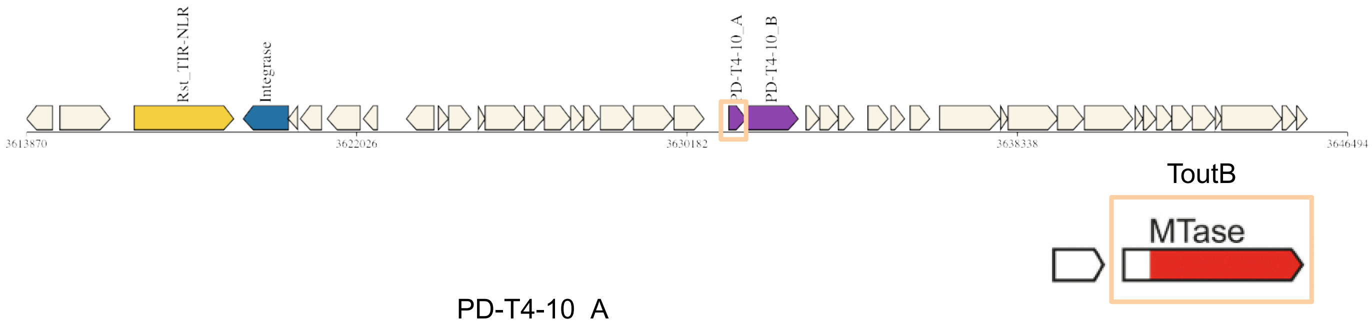


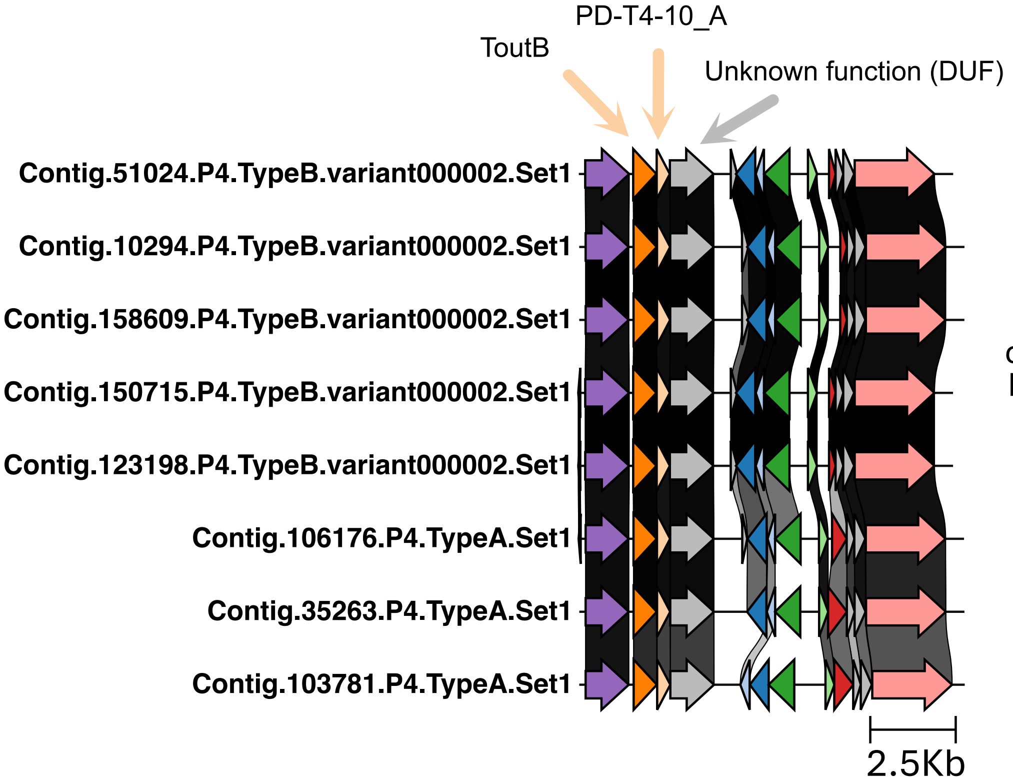


Another potential mix between systems, this one could also be two different systems that co-localize in P4. The locus consists of a Toutatis system, with only with toutB detected (but reported as complete by DefenseFinder), the PD-T4-10_A protein, and a third unknown protein.
