## Supplementary material for "Shuttling, swapping and mixing: the rapid modular evolution of antiviral repertoires in temperate phages and their satellites": FileS3_NovelSystemsAnnotations.pdf

### Crouga (Unknown 3A/3B) (pFD421)

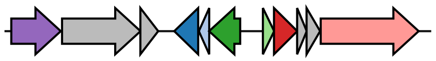

**Genomic Viewer** : Colocalization

*Notes:*
*RM-like system?*

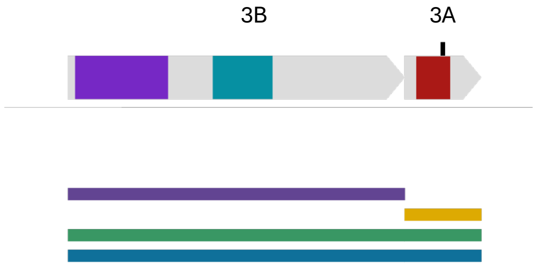


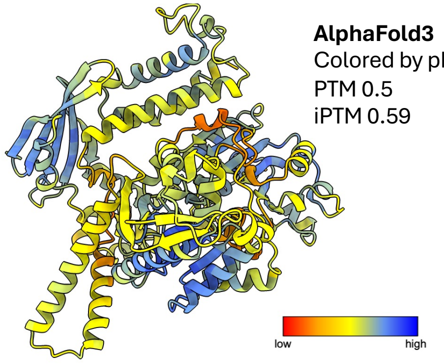


**AlphaFold3**
Colored by pLDDT score
PTM 0.5
iPTM 0.59

#### Unknown 3B: Nuclease/Helicase

**DefenseFinder Domains**: RM__Type_I_REases_FAM_2.einsi_trimmed

**GeneCLR hits**: VAPO_6407

**MGE**: 5 kb to Integrease / Satellite

**Pfam domains in homologs and original sequence:**

ResIII (PF04851)

DEAD (PF00270)

Helicase C (PF00271)

#### Unknown 3A: putative immune protein

**Defense Score**: 0.19

**GeneCLR hits**: VAPO_6407

**MGE**: 5 kb to Integrease / Satellite

**Pfam domains in original sequence:**

DUF6156 (PF19653)

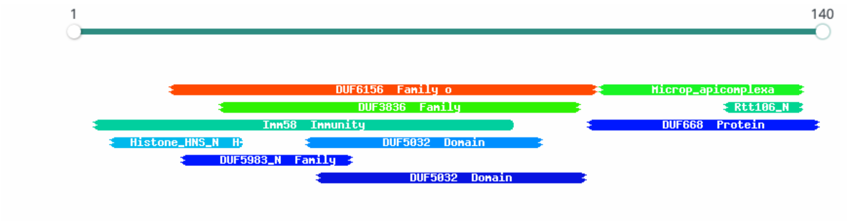

### Bandua (Unknown 4A/4B) (pFD422)

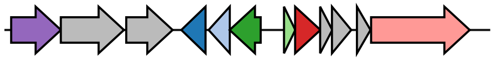

**Genomic Viewer :** Colocalization unsure

*Notes:*
*Unknown 4A has DS-23 signature*
*Unknown 4B has Stk2 signature*

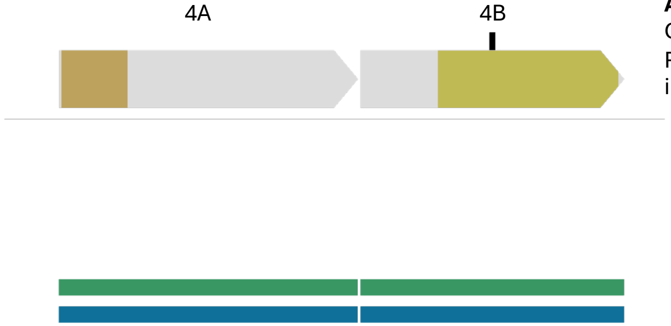


**AlphaFold3**
Colored by pLDDT score
PTM 0.59
iPTM 0.33

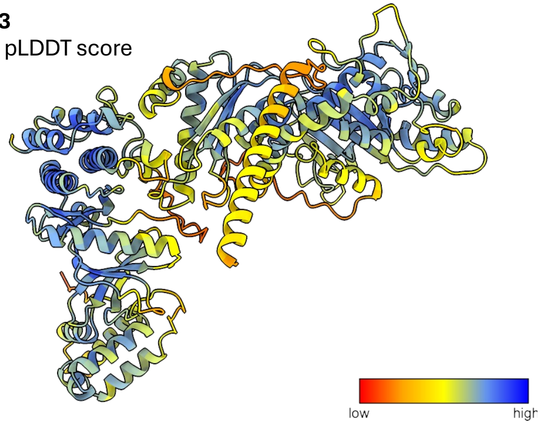


#### Unknown 4A: Putative nuclease

**GeneCLR hits**: PAPO_5975 or PAPP_367223

**MGE**: Satellite or 5 kb to Integrase

**Pfam domains in original sequence:**

Nuclease Related Domain – NERD / PDDEXK (PF08378)

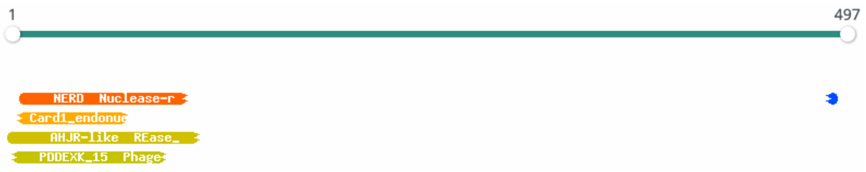


#### Unknown 4B: Kinase protein

**GeneCLR hits**: PAPP_420580 (no hit in a PAPO with Unknown4A)

**MGE**: Satellite or 5 kb to Integrase

**Pfam domains in homologs:**

PK_Tyr_Ser-Thr (PF07714)

**Pfam domains in original sequence:**

**HHpred hits**: PK_Tyr_Ser-Thr (PF07714)

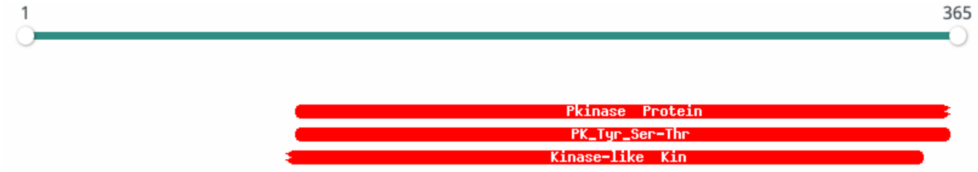

### Aernus (Unknown 5A/5B) (pFD423)

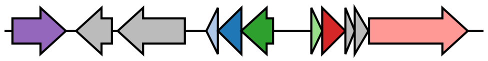

**Genomic Viewer** : Colocalization unsure

*Notes:*
*Unknown 5A PDDEXK class*

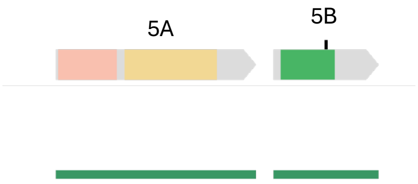


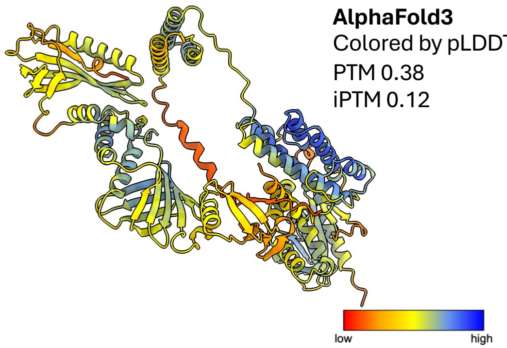


**AlphaFold3**
Colored by pLDDT score
PTM 0.38
iPTM 0.12

Unknown 5A: **Kinase/DNA binding protein/Nuclease**

**GeneCLR hits**: PAPO_8883 / PAPO_11039
when 5A is alone: PAPP_145658

**MGE**: 5 kb to Integrase or in Prophage when 5A alone

**Pfam domains in original sequence:**

DUF4365 (PF14280)

AbiGii (PF16873)

**FoldSeek top hits**: Holliday Junction Resolving Enzyme

**HHpred top hits**: PDDEXK (DUF4365) + AbiGii (PF16873)

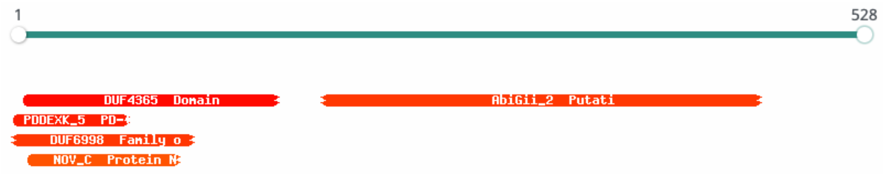


Unknown 5B: **Mannitol repressor**

**GeneCLR hits**: PAPO_8883 / PAPO_11039

**Pfam domains in homologs:**

Mannitol repressor *MtlR* (PF05068)

**Pfam domains in original sequence:**

**FoldSeek top hits**: MtlR-HPr complex

**HHpred top hits**: Mannitol repressor (PF05068)

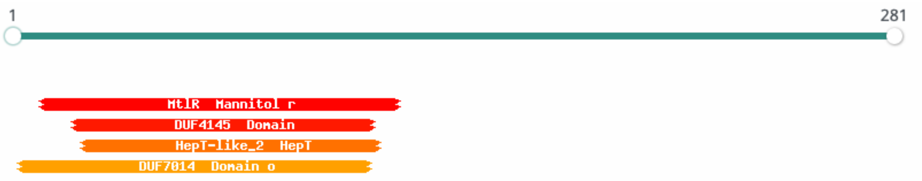

### Toga (Unknown 6: Gao_PpI-like / SMC-like) (pFD424)

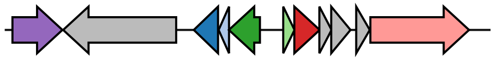

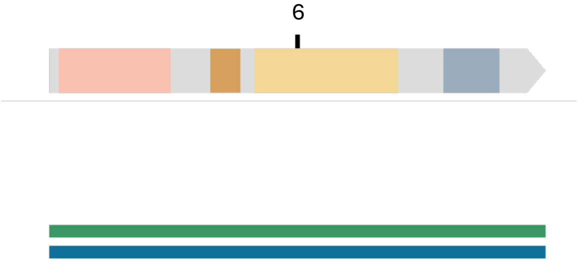


Unknown 6: **AAA_family_ATPase**

**AlphaFold3**
Colored by pLDDT score
PTM 0.84
iPTM NA

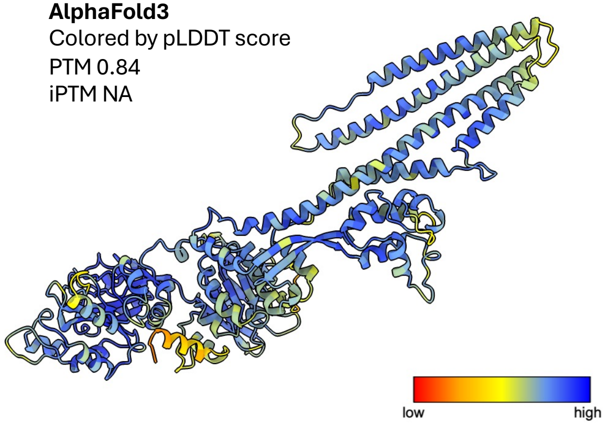


**GeneCLR hits**: PAPP_390102

**MGE**: Satellite

**Pfam domains in homologs:**

AAA_29 (PF13555)

**Pfam domains in original sequence:**

**FoldSeek top hits**: Smc head domain with a coiled coil and joint

CpsB_CapC (PF13555)

Macoilin (PF09726)

AAA_13 (PF13166)

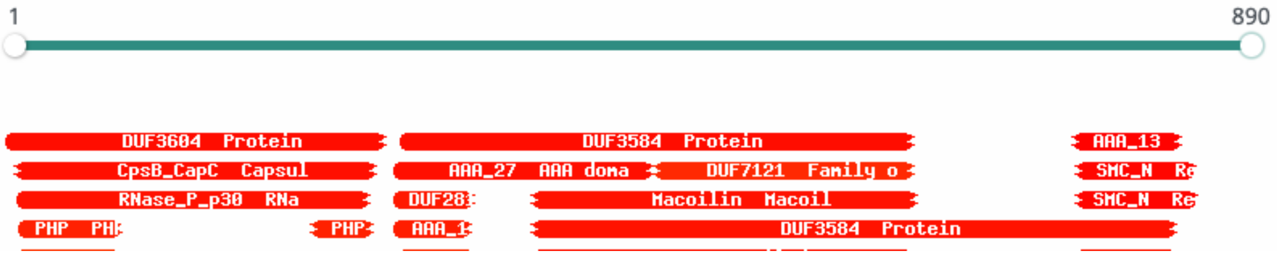

### Reve (Unknown 7: KAP-NTPase class) (pFD425)

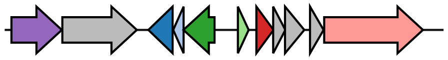

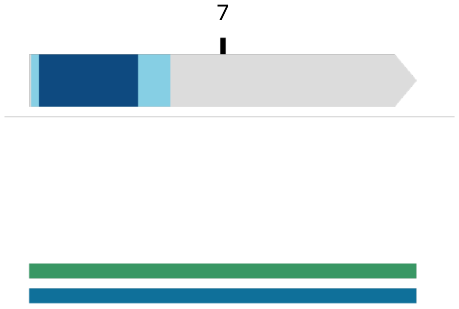


Unknown 7: **P-loop_NTPase_fold_protein**

**AlphaFold3**
Colored by pLDDT score
PTM 0.68
iPTM NA

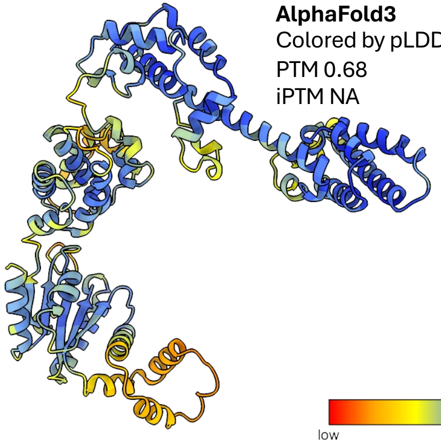

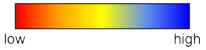


**GeneCLR hits**: PAPP_179762

**MGE**: Satellite

Pfam domains in homologs:

AAA domain (PF13401)

KAP family P-loop domain (PF07693)

Pfam domains in original sequence:

**FoldSeek top hits**: human Origin Recognition Complex ATPase motor module

**HHpred top hits**: KAP family P-loop domain (PF07693)

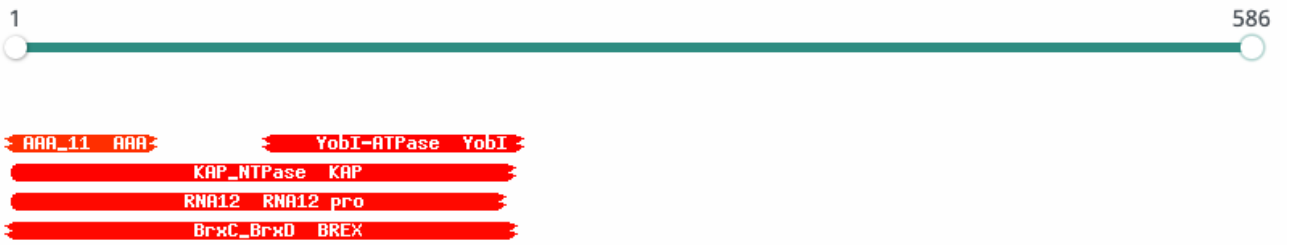

### Hailong (Unknown 8A/8B) (pFD426)

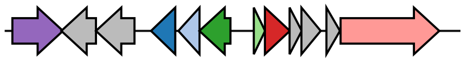

**Genomic Viewer** : Colocalization

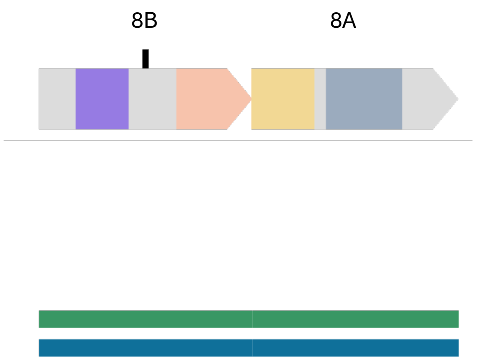


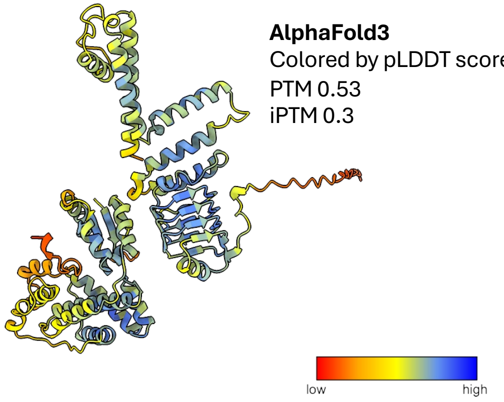


**AlphaFold3**
Colored by pLDDT score
PTM 0.53
iPTM 0.3

<u>Unknown 8B:</u>

**GeneCLR hits**: PAPP_180790

**MGE**: Satellite

**<u>Pfam domains in original sequence:</u>**

Pentapeptide_4 (PF13599)

IRK (PF01007)

**FoldSeek top hits**: Intermediates in the Gating of a K+ Channel

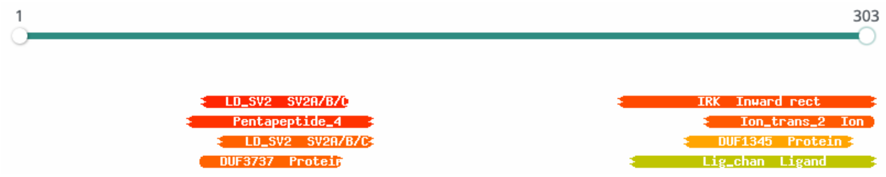


<u>Unknown 8A:</u>

**GeneCLR hits**: PAPP_180789

**MGE**: Satellite

**<u>Pfam domains in original sequence:</u>**

**FoldSeek top hits**: Kanamycin Nucleotidyltransferase

NTF-like (PF14540)

HEPN-like (PF18726)

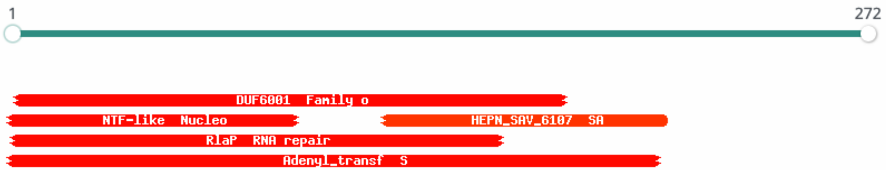
